# Cbfβ regulates Wnt/β-catenin, Hippo/Yap, and TGFβ signaling pathways in articular cartilage homeostasis and protects from ACLT surgery-induced osteoarthritis

**DOI:** 10.1101/2024.01.15.575763

**Authors:** Wei Chen, Yun Lu, Yan Zhang, Jinjin Wu, Abigail McVicar, Yilin Chen, Siyu Zhu, Guochun Zhu, You Lu, Jiayang Zhang, Matthew McConnell, Yi-Ping Li

**Author notes:** **Corresponding authors: Yi-Ping Li,** Department of Pathology and Laboratory Medicine, Tulane University School of Medicine, 1441 Canal St, Room 318, New Orleans, Louisiana, 70112, USA, **Wei Chen**, Department of Pathology and Laboratory Medicine, Tulane University School of Medicine, 1441 Canal St, Room 319, New Orleans, Louisiana, 70112, USA.

## Abstract

As the most common degenerative joint disease, osteoarthritis (OA) contributes significantly to pain and disability during aging. Several genes of interest involved in articular cartilage damage in OA have been identified. However, the direct causes of OA are poorly understood. Evaluating the public human RNA-seq dataset showed that *Cbfβ,* (subunit of a heterodimeric Cbfβ/Runx1,Runx2, or Runx3 complex) expression is decreased in the cartilage of patients with OA. Here, we found that the chondrocyte-specific deletion of *Cbfβ* in tamoxifen-induced *Cbfβ^f/f^Col2α1-CreER^T^* mice caused a spontaneous OA phenotype, worn articular cartilage, increased inflammation, and osteophytes. RNA-sequencing analysis showed that *Cbfβ* deficiency in articular cartilage resulted in reduced cartilage regeneration, increased canonical Wnt signaling and inflammatory response, and decreased Hippo/YAP signaling and TGF-β signaling. Immunostaining and western blot validated these RNA-seq analysis results. ACLT surgery-induced OA decreased *Cbfβ* and Yap expression and increased active β-catenin expression in articular cartilage, while local AAV-mediated *Cbfβ* overexpression promoted Yap expression and diminished active β-catenin expression in OA lesions. Remarkably, AAV-mediated *Cbfβ* overexpression in knee joints of mice with OA showed the significant protective effect of *Cbfβ* on articular cartilage in the ACLT OA mouse model. Overall, this study, using loss-of-function and gain-of-function approaches, uncovered that low expression of *Cbfβ* may be the cause of OA. Moreover, Local admission of *Cbfβ* may rescue and protect OA through decreasing Wnt/β-catenin signaling, and increasing Hippo/Yap signaling and TGFβ/Smad2/3 signaling in OA articular cartilage, indicating that local *Cbfβ* overexpression could be an effective strategy for treatment of OA.

## INTRODUCTION

As the most common degenerative joint disease, osteoarthritis (OA) is associated with painful, chronic inflammation that often leads to severe joint pain and joint stiffness for people over the age of 55(1, 2). Aging is a major contributor to OA, affecting the knees, hips, and spine and inflicting pain(1, 3–5). OA is characterized by a multitude of clinical and laboratory findings including osteophyte formation, cartilage degradation, subchondral bone thickening, and elevated cartilage degradation enzymes such as matrix metalloproteinases and aggrecanases (2, 6, 7). Treatment options for joint degeneration in OA are often palliative and oftentimes require surgical interventions such as joint replacement(8), but artificial joints can wear out or come loose and might eventually need to be replaced. As such, a more complete understanding of the mechanisms underlying how transcription factors regulate bone and cartilage formation to maintain bone and cartilage homeostasis could be critical to developing therapies for degenerative joint diseases such as OA.

Recent studies have begun to shed light on the nature of the genetic basis of OA and have confirmed several genes of interest involved in subchondral bone and articular cartilage degeneration including YAP, Sox9, Wnt/β-catenin signaling, and TGF-β/BMP signaling (4, 9–14). Core binding factors are heterodimeric transcription factors consisting of alpha (*Cbfα*) and beta (*Cbfβ*) subunits(15, 16). The *Cbfβ* subunit is a non-DNA-binding protein that binds *Cbfα* (also known as *Runx*) proteins to mediate the affinity of their DNA-binding (15, 16). *Runx/Cbfβ* heterodimers play critical roles in chondrocyte commitment, proliferation, and differentiation, as well as osteoblast differentiation (15–22). *Cbfβ* was reported as a potential key transcriptional factors in the regulatory network of OA by Gene Expression Omnibus data analysis(23). Yet the function of *Cbfβ* in OA pathogenesis remains unclear due to the lack of gain-of-function and loss-of-function animal model studies (15). Recently, another study has identified that *Cbfβ* may play an important role in regeneration and repair of articular cartilage in OA (24). Moreover, a recent study on a small molecule kartogenin showed the crucial role of *Cbfβ-Runx1* transcriptional program in chondrocyte differentiation in OA(25). However, the underlying mechanism behind *Cbfβ* regulation in OA remains unclear.

In this study, we showed that the deletion of *Cbfβ* in the postnatal cartilage in tamoxifen (TMX) induced *Cbfβ^f/f^Col2α1-CreER^T^* mice caused a spontaneous OA phenotype, including wear and loss of cartilage, osteophytes, decreased hip joint space, and increased inflammation. Notably, we observed the most severe phenotype in mutant mouse knee joints and hip joints. The loss-of-function study demonstrates the important role of *Cbfβ* in chondrocyte homeostasis and provides important insights into the role of *Cbfβ* as a critical transcriptional factor in OA. We also observed that *Cbfβ* enhanced articular cartilage regeneration and repair by modulating multiple key signaling pathways, including Hippo/YAP, Wnt/β-catenin, TGF-β, and Sox9. In addition, we demonstrated that adeno-associated virus-mediated local *Cbfβ* over-expression protects against surgery-induced OA in mice. The investigation of *Cbfβ*-multiple signaling regulation helps us better understand the OA genesis mechanism and will potentially facilitate the development of novel treatments for OA.

## RESULTS

### Tamoxifen (TMX) induced *Cbfβ^f/f^Col2α1-CreER^T^* developed spontaneous OA

To investigate the role of *Cbfβ* in spontaneous OA, the expression level of *Cbfβ* was first examined in human patients with OA by analyzing relevant datasets from published sources(26, 27) (**Fig. 1A, B**). Interestingly, there was significantly reduced *Cbfβ* gene expression in cartilage of human OA patients compared to healthy individuals (**Fig. 1A**)(26). Moreover, Methyl-seq data of human OA patient hip tissue exhibited increased methylation at the *Cbfβ* promoter of OA patients compared to healthy individuals, indicating inhibited *Cbfβ* expression in OA individuals may be through epigenetic regulation (**Fig. 1B**)(^27^). These data revealed that *Cbfβ* might play an important role in suppressing OA.

**Figure 1.**
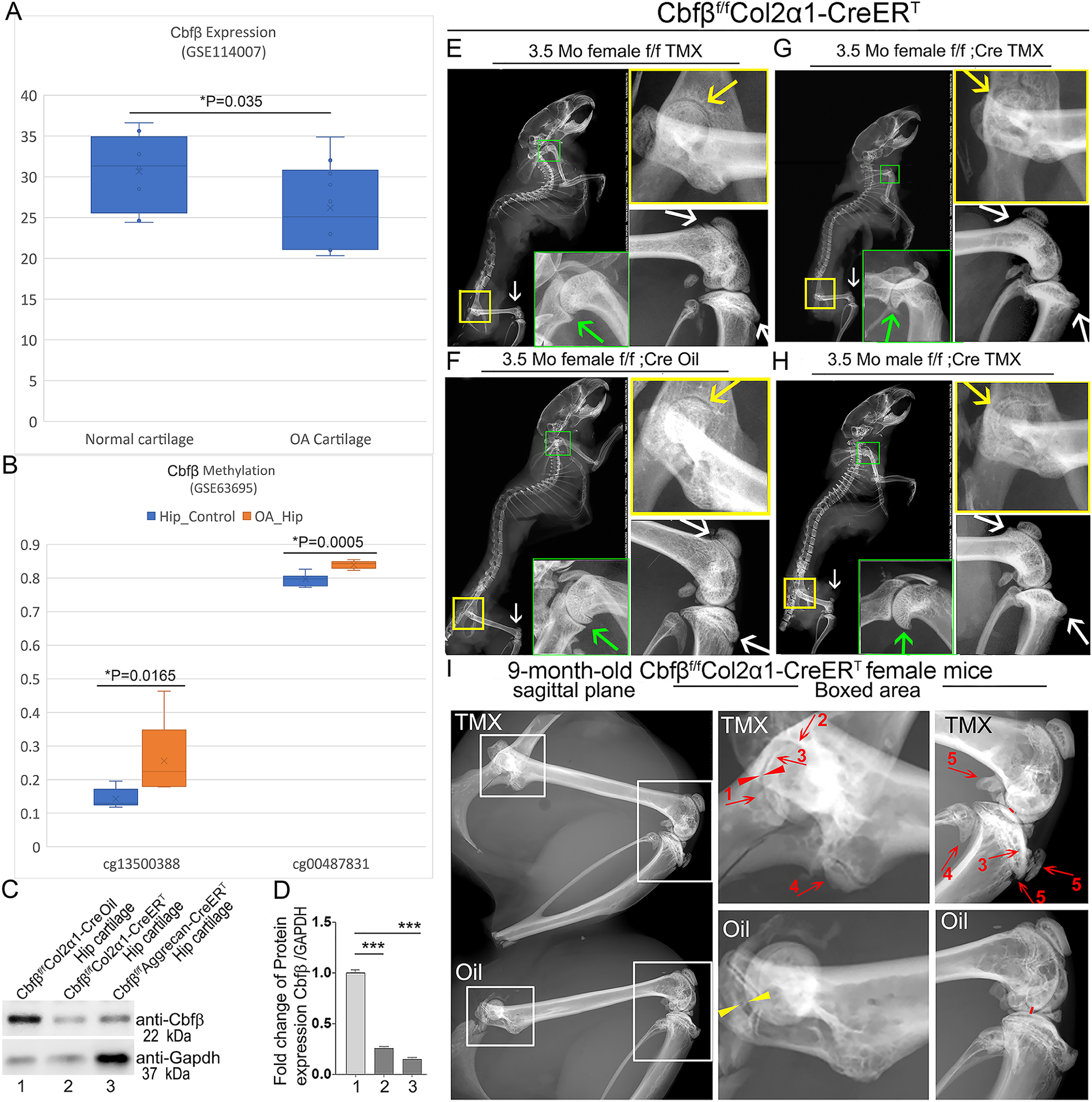
Tamoxifen (TMX) induced *Cbfβ^f/f^Col2α1-CreER^T^*mice developed spontaneous OA. **(A)** Public human RNA-seq dataset (n=8) (GSE114007) showing *Cbfβ* mRNA expression level in Normal and OA patient cartilage. (**B**) Public human methyl-seq dataset (n=5) (GSE63695) showing methylation at the *Cbfβ* promoter region (cg13500388 and cg00487831) in Normal and OA hip tissue. Statistical significance was assessed using Student’s t-test. Values were considered statistically significant at p<0.05. **(C)** Western blot to examine *Cbfβ* protein levels in the hip articular cartilage of 3.5-month-old male oil injected *Cbfβ^f/f^Col2α1-Cre* and TMX injected *Cbfβ^f/f^Col2α1-CreER^T^*, and 4-month-old male TMX injected *Cbfβ^f/f^Aggrecan-CreER^T^* mice (n=3). **(D)** Quantification of (C). **(E)** X-ray of 3.5-month-old TMX injected female *Cbfβ^f/f^* mouse hip, shoulder, and knee joint (n=15). **(F)** X-ray of 3.5-month-old oil injected female *Cbfβ^f/f^Col2α1- CreER^T^* mouse hip, shoulder, and knee joint (n=15). **(G)** X-ray of 3.5-month-old TMX injected female *Cbfβ^f/f^Col2α1-CreER^T^* mouse hip, shoulder, and knee joint (n=12). **(H)** X-ray of 3.5-month-old TMX injected male *Cbfβ^f/f^Col2α1-CreER^T^* mouse hip, shoulder, and knee joint. Green arrow: osteophytes in shoulder; yellow arrow: hip joint space; white arrow: hyperosteogeny in knee. **(I)** X-ray image of hips and knee joints of 9-month-old female *Cbfβ^f/f^Col2α1-CreER^T^* mice with oil injection and *Cbfβ^f/f^Col2α1-CreER^T^* mice with TMX injection (n=9). Red arrow 1,2,3: worn articular cartilage; Red arrow 4,5: osteophytes (spurs); Red arrow head: narrow joint space; Yellow arrow head: healthy hip joint space. **Figure 1-Source data.** Raw western blot images for Figure 1C.

Further, to evaluate the impact of *Cbfβ* loss-of-function on OA development, TMX inducible *Cbfβ^f/f^Col2α1-CreER^T^* and *Cbfβ^f/f^Aggrecan-CreER^T^* mice were generated by crossing *Cbfβ^f/f^* mice with either TMX inducible *Col2α1-CreER^T^*or *Aggrecan-CreER^T^* mouse lines. First, the validity of our mice models was confirmed by western blotting. *Cbfβ* protein levels were significantly decreased in the hip articular cartilage of both *Cbfβ^f/f^Col2α1-CreER^T^* and *Cbfβ^f/f^Aggrecan-CreER^T^* mice after TMX injection, indicating successful knockout of *Cbfβ* in both mouse models (**Fig. 1C, D**). Next, the bone phenotype in *Cbfβ* conditional knockout mice was examined. Whole-body X-ray images of 3.5-month-old male and female *Cbfβ^f/f^* and *Cbfβ^f/f^Col2α1-CreER^T^* mice after TMX injection showed osteophytes in the shoulder joint compared to *Cbfβ^f/f^Col2α1-CreER^T^*mice corn oil or *Cbfβ^f/f^* TMX injection controls (**Fig. 1E, G, F, H**, green arrows). X-ray results also revealed that in the TMX-induced *Cbfβ^f/f^Col2α1-CreER^T^* mice, the articular cartilage presented unclear borders and narrow hip joint spaces compared to the control groups (**Fig. 1E, G, F, H**, yellow arrows). *Cbfβ*-deficient mice also developed bone hyperosteogeny at the knee joints as shown by X-ray (**Fig. 1E, G, F, H**, white arrows). Moreover, TMX injected 9-month-old female *Cbfβ^f/f^Col2α1-CreER^T^* mice developed more severe OA phenotypes of joint blurred borders (worn articular cartilage red arrows 1, 2, and 3), osteophytes (bone spurs, red arrows 4 and5), and narrow joint space (red arrow heads) compared to the oil injected *Cbfβ^f/f^Col2α1-CreER^T^* controls (**Fig. 1I**, yellow arrow heads indicating healthy hip joint space). These data suggested that *Cbfβ*-deficient mice develop whole-body bone phenotypes that mimic human OA, and *Cbfβ* plays an important role in postnatal cartilage regeneration which affects OA onset and progression.

### Deficiency of *Cbfβ* in cartilage of 3.5-month-old mutant mice resulted in a more severe OA-like phenotype with decreased articular cartilage and osteoblasts, and increased osteoclasts and subchondral bone hyperplasia

To delve deeper into understanding the influence of *Cbfβ* in regulating the progression of OA, a chronological examination of hip joint histology was conducted, encompassing 1-month-old, 2-month-old, and 3.5-month-old TMX-induced *Cbfβ^f/f^Col2α1-CreER^T^* mice. Hematoxylin and eosin (H&E) and Safranin O (SO) staining of 1-month-old *Cbfβ^f/f^Col2α1-CreER^T^*mice (2 weeks after TMX induction) hip joints showed thicker femoral head cartilage (**Fig. 2A, B, E**, left panel) and slightly decreased tartrate-resistant acid phosphatase (TRAP)-positive cell numbers when compared to the controls (**Fig. 2C, F**, left panel). No significant change in alkaline phosphatase (ALP)-positive osteoblast numbers was detected (**Fig. 2D, G**, left panel). However, at 2 months old, *Cbfβ-*deficient mice (6 weeks after TMX induction) hip joints had about 2-fold cartilage loss in the femoral head (**Fig. 2A, B, E**, middle panel) with about 2-fold increased TRAP-positive osteoclast numbers, indicating increased inflammation (**Fig. 2C, F**, middle panel) and 3-fold decreased ALP-positive osteoblast numbers (**Fig. 2D, G**, middle panel). Additionally, a comparable pattern manifested in the hip joints of 3.5-month-old *Cbfβ^f/f^Col2α1-CreER^T^*mice (12 weeks after TMX induction). Notably, there were about 8-fold decrease in the SO-positive area (**Fig. 2A, B, E**, right panel), about 5.5-fold increase in TRAP-positive osteoclasts (**Fig. 2C, F**, right panel) and about 10-fold decrease in ALP-positive osteoblasts (**Fig. 2D, G**, right panel). It was noticed that there was significant subchondral bone hyperplasia in 3.5-month-old mutant mice (**Fig. 2C**, right panel). Collectively, histological data provided additional support, indicating that while *Cbfβ* did not exert a significant effect on the hip cartilage of 1 month-old mice, deficiency of Cbfβ in cartilage in 3.5-month-old mutant mice resulted in a more severe OA-like phenotype with decreased articular cartilage and osteoblasts, and increased osteoclasts and subchondral bone hyperplasia. Our data further supported that *Cbfβ* plays a crucial role in articular cartilage regeneration, and deficiency of *Cbfβ* in mice might lead to the progression of OA.

**Figure 2.**
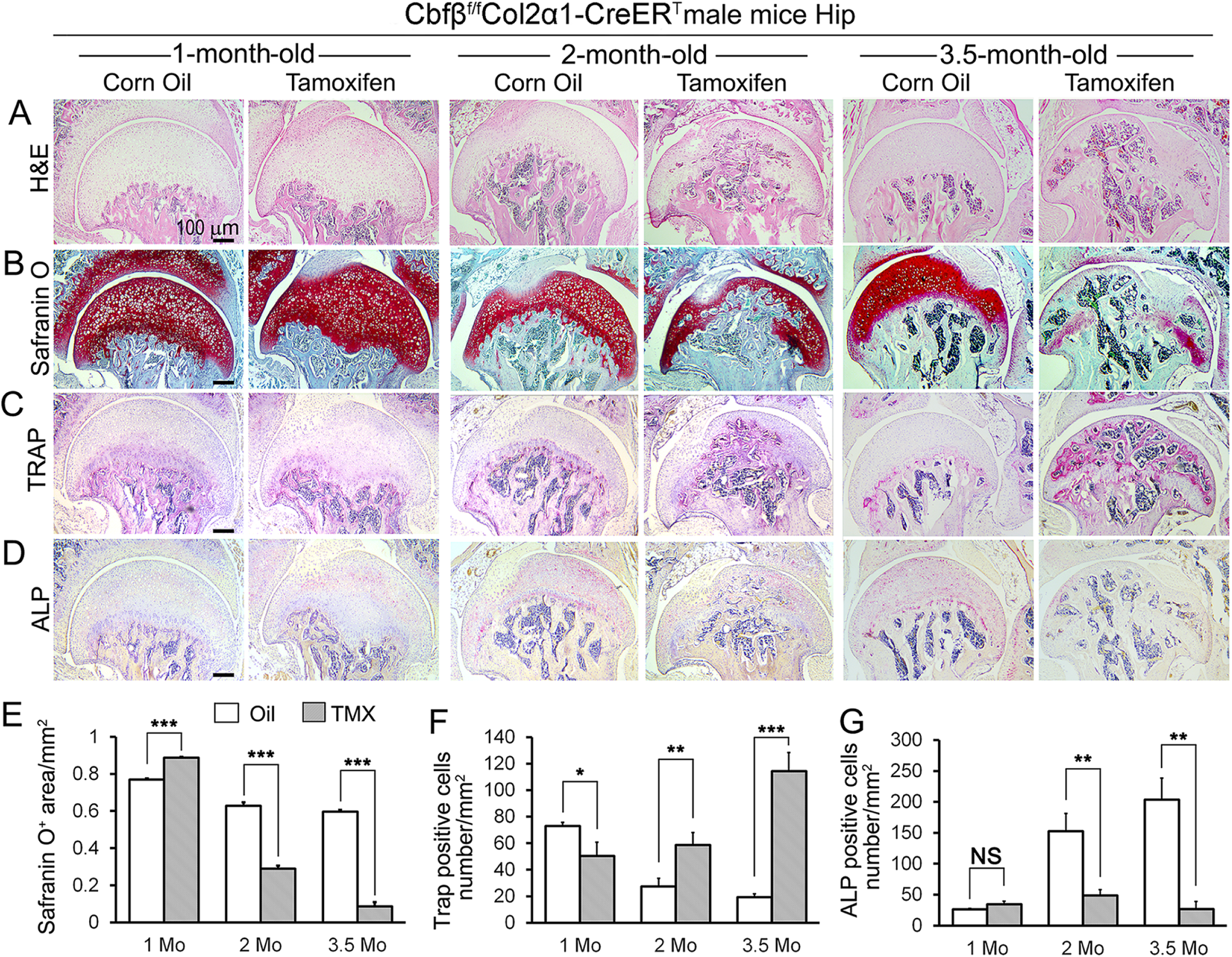
*Cbfβ* deletion in *Col2α1-CreER^T^* mice cartilage resulted in more severe OA-like phenotype 3.5-month-old mutant mice with increased osteoclasts and subchondral bone hyperplasia, decreased articular cartilage and osteoblasts. **(A-D)**. H&E staining **(A)**, SO staining **(B)**, TRAP staining **(C)**, and ALP staining **(D)** of 1-month-old, 2-month-old, and 3.5-month-old male *Cbfβ^f/f^Col2α1-CreER^T^* mice hips respectively. **(E)** Quantification of SO red area of (B). Data was measured by ImageJ. **(F)** Quantification of TRAP-positive cell numbers of (C). **(G)** Quantification of ALP-positive cell numbers of (D). TMX=Tamoxifen, *Cbfβ* deleted group; Oil=Corn Oil, control group. n=7. Data are shown as mean ± SD. NS, no significance; *p<0.05; **p<0.01; ***p<0.001 vs. controls by Student’s t-test. Scale bar: 100μm.

### The deficiency of *Cbfβ* may be the cause of early onset OA

Anterior cruciate ligament (ACL) injury is a common cause of human OA, and Anterior cruciate ligament transection (ACLT) is a well-established mouse model that mimics human OA. Bone remodeling between chondrocytes and subchondral bone ossification is known to be important for OA(7). In order to further analyze the role of *Cbfβ* in OA pathological conditions, we developed OA pathological disease mice models by performing ACLT surgery on mice knees. Then, we performed radiographical and histological studies on WT, *Cbfb^f/f^*, and *Cbfβ^f/f^Col2α1-CreER^T^* mice with or without ACLT surgery. We discovered that in *Cbfβ^f/f^Col2α1-CreER^T^* mice with ACLT surgery, more severe articular cartilage wear (white arrow) showed unclear borders, joint space loss (purple arrow), more hyperosteogeny (blue arrow), and significantly enhanced subchondral bone density (red arrow) compared to the control groups, indicating spontaneous OA-like symptoms (**Fig. 3A, B**). Moreover, *Cbfβ^f/f^Col2α1-CreER^T^*mice with no ACLT surgery knee joint space has narrower joint space compared to WT mice with ACLT surgery (Red Arrowhead) (**Fig. 3A, B**). Those results show that *Cbfβ* deficiency accelerated the development of OA in the *Cbfβ^f/f^Col2α1-CreER^T^*mice with ACLT surgery. Moreover, SO staining also showed that *Cbfβ^f/f^Col2α1-CreER^T^* mice with ACLT surgery had less SO-positive area compared to *Cbfβ^f/f^* mice with ACLT surgery, indicating increased cartilage loss (**Fig. 3C, D**). The Osteoarthritis Research Society International (OARSI) Score analysis showed that *Cbfβ^f/f^Col2α1-CreER^T^*TMX injected mice with no surgery presented similar OARSI Score compared with *Cbfb^f/f^*mice with ACLT surgery, indicating the important role of *Cbfβ* in articular cartilage homeostasis (**Fig. 3E).** Interestingly, *Cbfβ^f/f^Col2α1-CreER^T^* TMX mice with ACLT surgery had a significantly increased OARSI Score compared to *Cbfβ^f/f^Col2α1-CreER^T^*TMX injected mice with no surgery and *Cbfb^f/f^* mice with ACLT surgery (**Fig. 3E).** Those results indicate that *Cbfβ* also plays an important role in regulating postnatal cartilage regeneration as well as bone destruction in OA pathological condition, and demonstrated that the deficiency of *Cbfβ could* be the cause of early onset OA.

**Figure 3.**
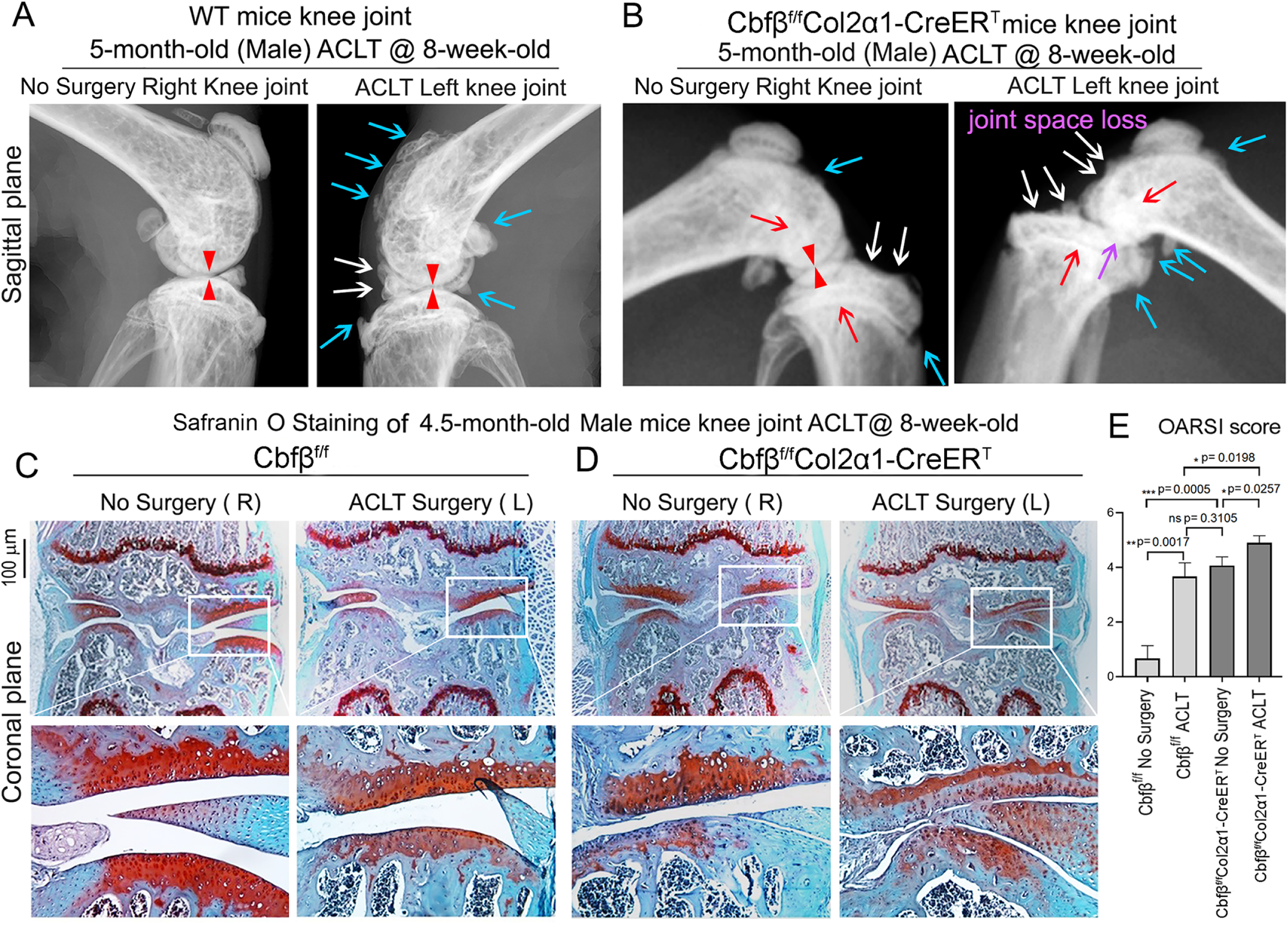
*Cbfβ^f/f^Col2α1-CreER^T^*mice with ACLT surgery developed early onset OA. **(A)** X-ray of 5-month-old male WT (ACLT at 8-weeks-old) mice knees (n=15). **(B)** X-ray of 5-month-old male *Cbfβ^f/f^Col2α1-CreER^T^* (ACLT at 8-weeks-old) mice knees. Red arrows indicate subchondral bone; Red arrow heads indicate joint space; Light blue arrows indicate osteophytes; White arrows indicate worn articular cartilage; Purple arrow indicates joint space loss; (n=15). **(C)** SO stain of 4.5-month-old male *Cbfβ^f/f^* (ACLT at 8-weeks-old) mice knees (n=7). **(D)** SO stain of 4.5-month-old male *Cbfβ^f/f^Col2α1-CreER^T^*(ACLT at 8-weeks-old) mice knees (n=6). **(E)** Knee joint Osteoarthritis Research Society International (OARSI) score of (C) and (D). Data are shown as mean ± SD. Scale bar: 100μm (**C-D**).

### RNA-seq analysis indicated that deficiency of Cbfβ in cartilage reduces cell fate commitment, cartilage regeneration and repair, and increases canonical Wnt signaling and inflammatory response

To dissect the mechanism underlying the role of *Cbfβ* in the articular cartilage regeneration in OA, genome-wide RNA-sequencing analysis was conducted using hip articular cartilage of 2-month-old *Cbfβ^f/f^Col2α1-CreER^T^* TMX injected mice compared with 2-month-old WT mice (**Fig. 4**). Volcano plot results illustrated that the top downregulated genes included Fabp3, Nmrk2, Csf3r, Rgs9, Plin5, Rn7sk, and Eif3j2 while top upregulated genes included Cyp2e1, Slc15a2, Alas2, Hba-a2, Lyve1, Snca, Serpina1b, Hbb-b1, Rsad2, Retn, and Trim10 in the articular cartilage of *Cbfβ* conditional knockout mice (**Fig. 4A**). Pie chart of articular cartilage from *Cbfβ^f/f^Col2α1-CreER^T^*mice demonstrated significantly altered differentially expressed genes (DEGs), where 70.7% were upregulated and 29.3% were downregulated (**Fig. 4B**). Among them, Rsad2 is known to be closely related to immune regulation and play a role in driving the inflammatory response through the NF-κB and JAK-STAT pathways(28). Increased expression of Rsad2 indicates that *Cbfβ* conditional knockout is associated with increased inflammatory signaling in mice knee joints (**Fig. 4A**). Moreover, several genes related to lipid metabolism and transport were downregulated in response to Cbfβ conditional knockout (**Fig. 4A**). Fabp3 is known to be involved in several processes, including lipid homeostasis and transport, and positive regulation of long-chain fatty acid import into cell (29, 30). In addition, Plin5 is a negative regulator of peroxisome proliferator activated receptor (PPAR) signaling, a positive regulatory of sequestering of triglyceride and regulation of lipid metabolic process(31). A previous study has shown that dysregulated lipid content or metabolism in cartilage leads to dysfunction cartilage(32). Decreased expression of Fabp3 and Plin5 in Cbfβ conditional knockout mice indicates the important positive regulatory role of Cbfβ in lipid transport and metabolism in articular cartilage, which is important in cartilage homeostasis (**Fig. 4A**).

**Figure 4.**
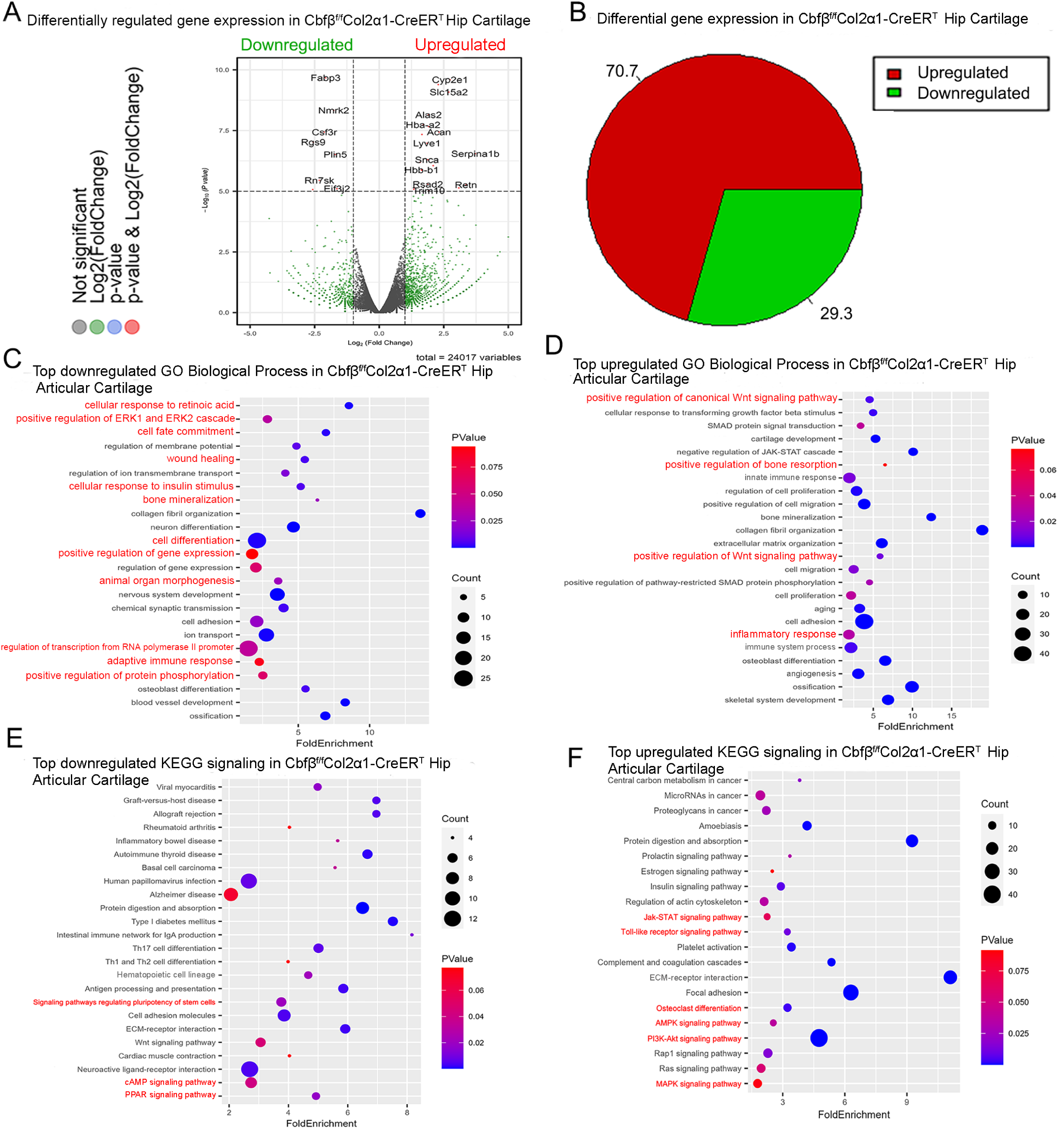
RNAseq analysis indicated that deficiency of Cbfβ in cartilage reduces cell fate commitment, cartilage regeneration and repair, and increases canonical Wnt signaling and inflammatory response. **(A)** Volcano plot showing differentially regulated gene expression in 6-weeks-old male *Cbfβ^f/f^* and *Cbfβ^f/f^Col2α1-CreER^T^* mice hip articular cartilage. **(B)** Pie chart showing percentage of upregulated and downregulated differentially regulated genes in hip articular cartilage of 6-weeks-old male *Cbfβ^f/f^Col2α1-CreER^T^*mice compared to those of *Cbfβ^f/f^* mice. The percentages of genes upregulated and downregulated are shown in red and green, respectively. **(C)** GO functional clustering of the top downregulated biological process (BP) in 6-week-old male *Cbfβ^f/f^Col2α1-CreER^T^*mice hip articular cartilage. **(D)** GO functional clustering of the top upregulated BP in 6-week-old male *Cbfβ^f/f^Col2α1-CreER^T^* mice hip articular cartilage. **(E)** GO functional clustering of the top downregulated KEGG signaling pathways in 6-week-old male *Cbfβ^f/f^Col2α1-CreER^T^*mice hip articular cartilage. **(F)** GO functional clustering of the top upregulated KEGG signaling pathways in 6-week-old male *Cbfβ^f/f^Col2α1-CreER^T^*mice hip articular cartilage.

To further investigate the functions of the differential expressed genes in *Cbfβ* conditional knockout mice, Gene Ontology (GO) studies were performed on both the upregulated and downregulated DEGs in *Cbfβ^f/f^Col2α1-CreER^T^* mice TMX injected compared to WT mice (**Fig 4C-F**). GO annotation based on GO Biological Processes (BP) showed significantly downregulated differentially expressed gene groups associated with Cellular Response to Retinoic Acid, Wound Healing, positive Regulation of Protein Phosphorylation in response to *Cbfβ* conditional knockout in *Cbfβ^f/f^Col2α1-CreER^T^* mice, further supporting the important role of *Cbfβ* in cartilage and bone development (**Fig. 4C**). Moreover, it was previously reported that p38/ERK/JNK/SMAD pathways are crucial in the chondrogenic differentiation induced by TGF-β1(33). GO BP analysis also revealed significantly downregulated genes in positive regulation of ERK1 and ERK2 cascade in *Cbfβ^f/f^Col2α1-CreER^T^* mice, indicating that *Cbfβ* deficiency in chondrocytes was associated with downregulated ERK signaling which resulted in dysregulated chondrocyte differentiation (**Fig. 4C**). Furthermore, *Cbfβ* conditional knockout is also associated with downregulated cell fate commitment, cell differentiation, positive regulation of gene expression, animal organ morphogenesis, regulation of RNA polymerase II promoter, and positive regulation of protein phosphorylation (**Fig. 4C**). Enrichment analysis of downregulated KEGG signaling pathways also demonstrated that *Cbfβ* deficiency in *Cbfβ^f/f^Col2α1-CreER^T^*mice led to significant changes in signaling pathways regulating pluripotency of stem cells (**Fig. 4E**). These results implied that *Cbfβ* deficiency leads to downregulated chondrocyte differentiation and proliferation. In addition, downregulated differential expressed genes in *Cbfβ* deficient mice were associated with cellular response to insulin stimulus (**Fig. 4C**). Previous studies have shown that insulin has anti-inflammatory effect by negatively regulating NF-κB, PI3k/AKT, and TLR signaling *etc.* (34, 35). Downregulated cellular response to insulin stimulus in *Cbfβ^f/f^Col2α1-CreER^T^* mice hip articular cartilage suggested dysregulated and elevated immune signaling in *Cbfβ* deficient mice articular cartilage. In addition, a recent study has shown that the activated cAMP pathway inhibits OA development(36). cAMP signaling is downregulated in Cbfβ deficient mice, indicating that Cbfβ may also regulate cAMP signaling in OA pathogenesis (**Fig. 4E**). On the other hand, top downregulated GO KEGG analysis in Cbfβ deficient cartilage also shows downregulated PPAR signaling, in line with decreased Plin5 expression shown in the volcano plot (**Fig. 4A, E**). As mentioned previously, PPAR signaling is crucial for cell differentiation and lipid metabolism(37, 38). Downregulated PPAR signaling indicated a crucial role of Cbfβ in articular cartilage regeneration and regulation of lipid content.

Upregulated GO BP and GO KEGG analysis results further elucidated the regulatory mechanism of *Cbfβ* in mice articular cartilage (**Fig. 4**). Firstly, upregulated GO BP pathways displayed significantly upregulated differentially expressed genes in the positive regulation of the canonical Wnt signaling pathway, indicating that *Cbfβ* negatively regulated canonical Wnt signaling pathway in articular cartilage (**Fig. 4D**). Besides, upregulated positive regulation of bone resorption in *Cbfβ* conditional knockout mice supported the bone destruction seen in previous phenotypical studies, showing the crucial role of *Cbfβ* in protecting against bone destruction (**Fig. 4D**). Further, both GO BP and GO KEGG results unveiled upregulated signaling pathways related to inflammatory response (**Fig. 4D, F**). Upregulated GO BP pathways including innate immune response, inflammatory response, immune system process, and angiogenesis were associated with *Cbfβ* deficiency and are also related to inflammation (**Fig. 4D**). Furthermore, enrichment analysis of upregulated KEGG signaling pathways demonstrated that *Cbfβ* deficiency led to significant changes in the JAK-STAT signaling pathway, Toll-like receptor signaling pathway, AMPK signaling pathway, and MAPK signaling pathway (**Fig. 4F**). The JAK/STAT pathway played an important role in multiple crucial cellular processes such as the induction of the expression of some key mediators that were related to cancer and inflammation(39). Moreover, studies had demonstrated the upregulation of TLR signaling in osteoarthritis (OA), highlighting its involvement in the induction of chondrocyte apoptosis (40), along with the pivotal role played by MAPK signaling in the pathogenesis of OA (41). These GO data indicated an augmentation in the positive regulation of the JAK-STAT cascade, TLR signaling, and MAPK signaling pathways following *Cbfβ* deletion, suggesting that the deficiency of *Cbfβ* led to an intensification of immune signaling contributing to the progression of osteoarthritic pathological processes. In addition, the downregulation of the adaptive immune response and the upregulation of the innate immune response further demonstrated that *Cbfβ* deficiency in the knee joint of mice was associated with heightened innate immune signaling while concurrently dampening adaptive immune signaling (**Fig 4C, D**).

### Heatmap analysis uncovered that *Cbfβ* deficiency in cartilage resulted in decreased chondrocyte genes expression and decreased TGF-β and Hippo signaling, but increased Wnt signaling

To further uncover the regulatory mechanism by which *Cbfβ* initiates signaling pathway changes in OA at the individual gene level, the gene expression profiles associated with chondrocytes, as well as with the Hippo, TGF-β, and Wnt signaling pathways were examined (**Fig. 5**). Given that OA is a systemic joint disease, an analysis was conducted on both the articular cartilage of the hip joint in *Cbfβ^f/f^Col2α1-CreER^T^* mice and the articular cartilage of the knee joint in *Cbfβ^f/f^Aggrecan-CreER^T^*mice (**Fig. 5**). Interestingly, chondrocyte-related genes were downregulated in the hip joint articular cartilage of the *Cbfβ^f/f^Col2α1-CreER^T^* mice. Other downregulated genes in the *Cbfβ*-deficient mice articular cartilage included Bmp2 and Runx2 (**Fig. 5A**). The *Cbfβ* subunit is a non-DNA-binding protein that binds *Cbfα* (also known as *Runx*) proteins to mediate their DNA-binding affinities. *Runx/Cbfβ* heterodimers play key roles in various developmental processes (15–22). Moreover, Bmp2 is a crucial protein in the development of bone and cartilage, a central protein in TGFβ signaling, and some of its specific functions include activating osteogenic genes such as Runx2(42). The downregulation of Bmp2 and Runx2 in *Cbfβ^f/f^Col2α1-CreER^T^* hip articular cartilage suggested an important role of *Cbfβ* in regulating articular cartilage generation through TGFβ signaling. Many genes were also upregulated in the *Cbfβ*-deficient articular cartilage, such as Adamts12 and Fgf2 (**Fig. 5A**). High expression of Adamts is a typical feature of OA, implying that *Cbfβ* deficiency may control the expression of Adamts to affect the differentiation of chondrocytes. Further, Fgf2 is previously reported to activate Runx2 via MER/ERK signaling pathway and increase MMP13 expression(43). Increased expression of Fgf2 was seen in both *Cbfβ^f/f^Col2α1-CreER^T^*hip articular cartilage as well as *Cbfβ^f/f^Aggrecan-CreER^T^* knee cartilage, showing *Cbfβ* might upregulate MAPK/ERK signaling in cartilage through Fgf2 (**Fig 5A**).

**Figure 5.**
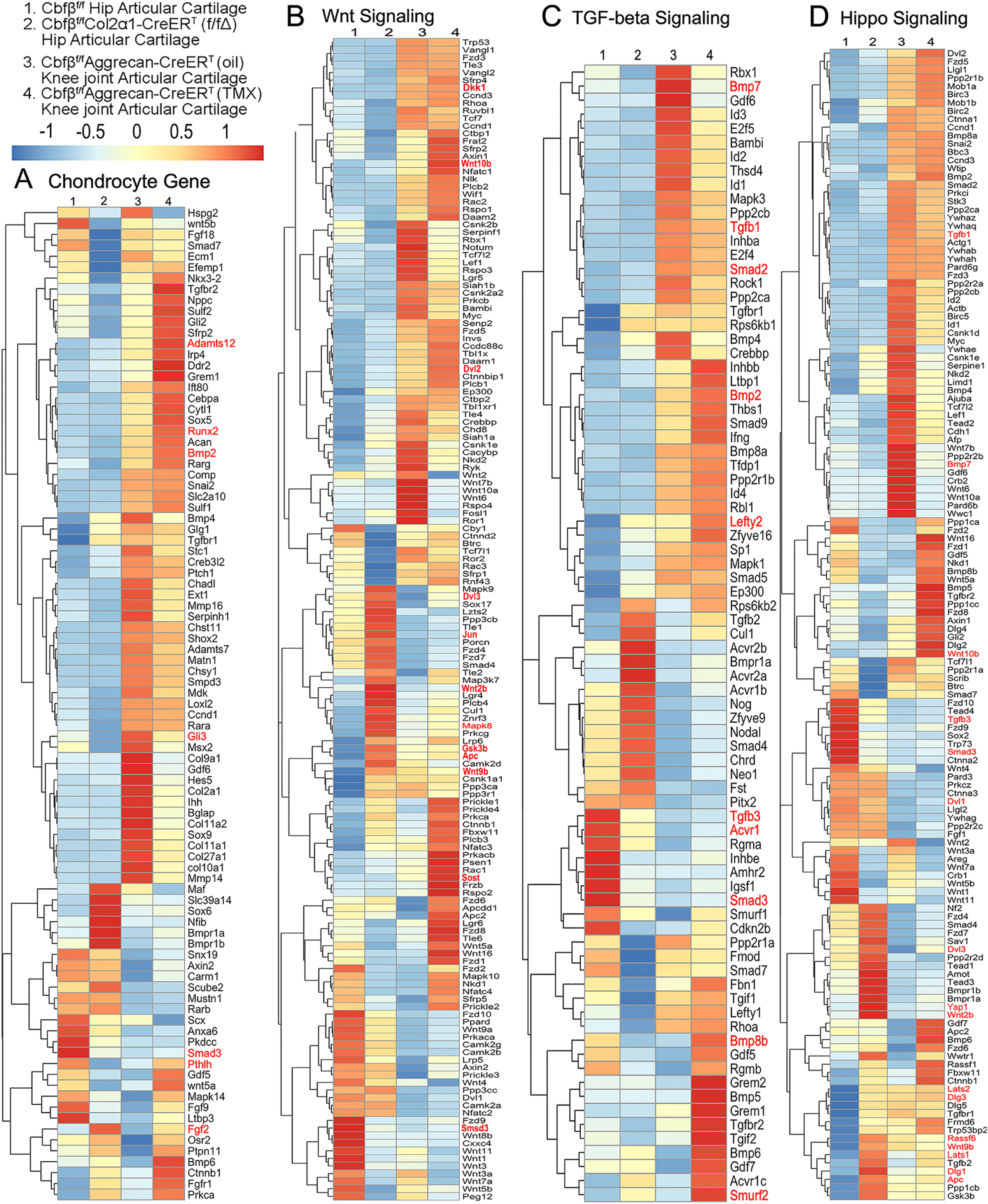
Heatmap analysis uncovered that deficiency of *Cbfβ* in cartilage resulted in decreased chondrocyte genes expression and decreased TGF-β and Hippo signaling, but increased Wnt signaling. **(A)** Heatmap for chondrocyte gene expression in (1) 6-weeks-old male *Cbfβ^f/f^* mice hip articular cartilage, (2) 6-weeks-old male *Cbfβ^f/f^Col2α1-CreER^T^*mice hip articular cartilage, (3) 12-weeks-old male oil injected *Cbfβ^f/f^Aggrecan-CreER^T^* mice knee joint articular cartilage, and (4) 12-weeks-old male *Cbfβ^f/f^Aggrecan-CreER^T^* mice (TMX injected at 6-weeks-old) knee joint articular cartilage. **(B)** Heatmap showing Wnt signaling-related gene expression. **(C)** Heatmap showing TGF-β signaling-related gene expression. **(D)** Heatmap showing Hippo signaling-related gene expression.

Moreover, the heatmap of RNA-seq analysis showed that *Cbfβ^f/f^Col2α1-CreER^T^* cartilage had altered gene expression levels in the Wnt, TGF-β, and Hippo signaling pathways (**Fig. 5**). Our results demonstrated that genes associated with Wnt signaling pathway activation, such as Mapk8, Dvl3, Wnt10b, Wnt2b, Wnt9b, and Jun(44) were upregulated, while the inhibitor of the Wnt signaling pathway Sost was downregulated, indicating that loss of *Cbfβ* could promote cartilage ossification and osteophyte formation through its activation of the Wnt pathway (**Fig. 5B**). Dvl3 is a positive regulator of the Wnt/β-catenin pathway, which can stabilize β-catenin and upregulate downstream target genes by interacting with Mex3a(45). These results suggested that the loss of *Cbfβ* could promote the expression of the activator of Wnt signaling, resulting in the activation of the Wnt signaling pathway.

Furthermore, our results also exemplified that the TGF-β signaling pathway repressors Lefty2 and Smurf2 were upregulated in the *Cbfβ*-deficient articular cartilage(46) (**Fig. 5C**). In addition, other genes involved in TGF-β signaling, such as Tgfb1, Acvrl1, Bmp7, Smad2, and Smad3, were downregulated in the *Cbfβ*-deficient articular cartilage (**Fig. 5C**). These results demonstrate that loss of *Cbfβ* leads to decreased expression of genes in TGF-β signaling and increased expression of repressors of TGF-β signaling, which results in the inhibition of the TGF-β signaling pathway.

Genes involved in the canonical Hippo signaling pathway such as APC, Dlg1, and Dlg3 were upregulated, signifying a close relationship of *Cbfβ* to Hippo signaling(**Fig. 5D**). APC is the downstream part of the Wnt signaling pathway, and through the cross-talk of Wnt signal and Hippo signal, APC mutation leads to the nuclear localization of YAP/TAZ and activates YAP-Tead and TAZ-Tead dependent transcription, and ultimately, Hippo signal is turned off(47). In our study, APC expression was enhanced in *Cbfβ*-deficient articular cartilage, supporting that *Cbfβ* deficiency in articular cartilage affected Hippo signaling (**Fig. 5D**). Lats1 and Lats2 are essential components of the Hippo pathway that phosphorylate and inactivate YAP, which is a key link in the activation and shutdown of the Hippo signaling pathway(48). Our study demonstrated that Lats1/Lats2 expression was enhanced in *Cbfβ*-deficient articular cartilage (**Fig. 5D**). Therefore, although there is increased Yap1 gene expression in *Cbfβ*-deficient mice, upregulated Lats1/2 potentially leads to increased phosphorylation in Yap protein and activated Hippo signaling pathway (**Fig. 5D**). Thus, loss of *Cbfβ* could inhibit the repressor of the Hippo signaling pathway and promote the expression of the activator of Hippo signaling, resulting in the activation of the Hippo signaling pathway. Examination of the expression profiles of these genes showed altered expression between the mutant and WT samples, with different expression patterns between *Cbfβ*-deficient articular cartilage in mice hip samples and *Cbfβ*-deficient knee samples, indicating that *Cbfβ* regulation is tissue-specific (**Fig. 5A-D**). Collectively, we are the first to demonstrate that *Cbfβ* may control downstream gene expression by orchestrating the TGF-β, Hippo, and Wnt signaling pathways, thereby setting off the cascade of OA pathological processes, including cartilage damage and inflammation.

### Postnatal *Cbfβ* deficiency in cartilage resulted in increased Wnt signaling, inflammatory genes expression, decreased cartilage formation genes expression in the knee articulate cartilage

To further investigate OA related genes expression of C*bfβ* deficiency mice in articular cartilage in which *Cbfβ* regulates articular cartilage regeneration, we performed immunohistochemistry (IHC) staining on *Cbfβ*-deficient mouse hip joints. The result showed that postnatal *Cbfβ* deficiency in cartilage (**Fig. 6A, F**) resulted in increased inflammatory genes expression, including decreased cartilage formation genes expression in the knee articulate cartilage. The chondrocytes cell markers Col2α1, and cartilage degradation markers MMP13 and ADAMTS5 were examined by IHC staining (**Fig. 6B, C, D, G**). As expected, mutant mice articular cartilage had significant degradation with low expression of Col2α1 in both the superficial zone and the deep zone, and the middle zone was replaced by bone with no Col2α1 expression (**Fig. 6B, G**). Aggrecanases (ADAMTSs) and matrix metalloproteinases (MMPs), especially ADAMTS5 and MMP13 are known to have important roles in cartilage destruction in OA. IHC staining results show without Cbfβ, articular cartilage had high expression of ADAMTS5 (**Fig. 6C, G**) and MMP13 (**Fig. 6D,G**), indicating mutant mice cartilage was undergoing severe cartilage degradation and increased inflammation. Negative control of the IHC staining shows the validity of the experiment (**Fig. 6E**).

**Figure 6.**
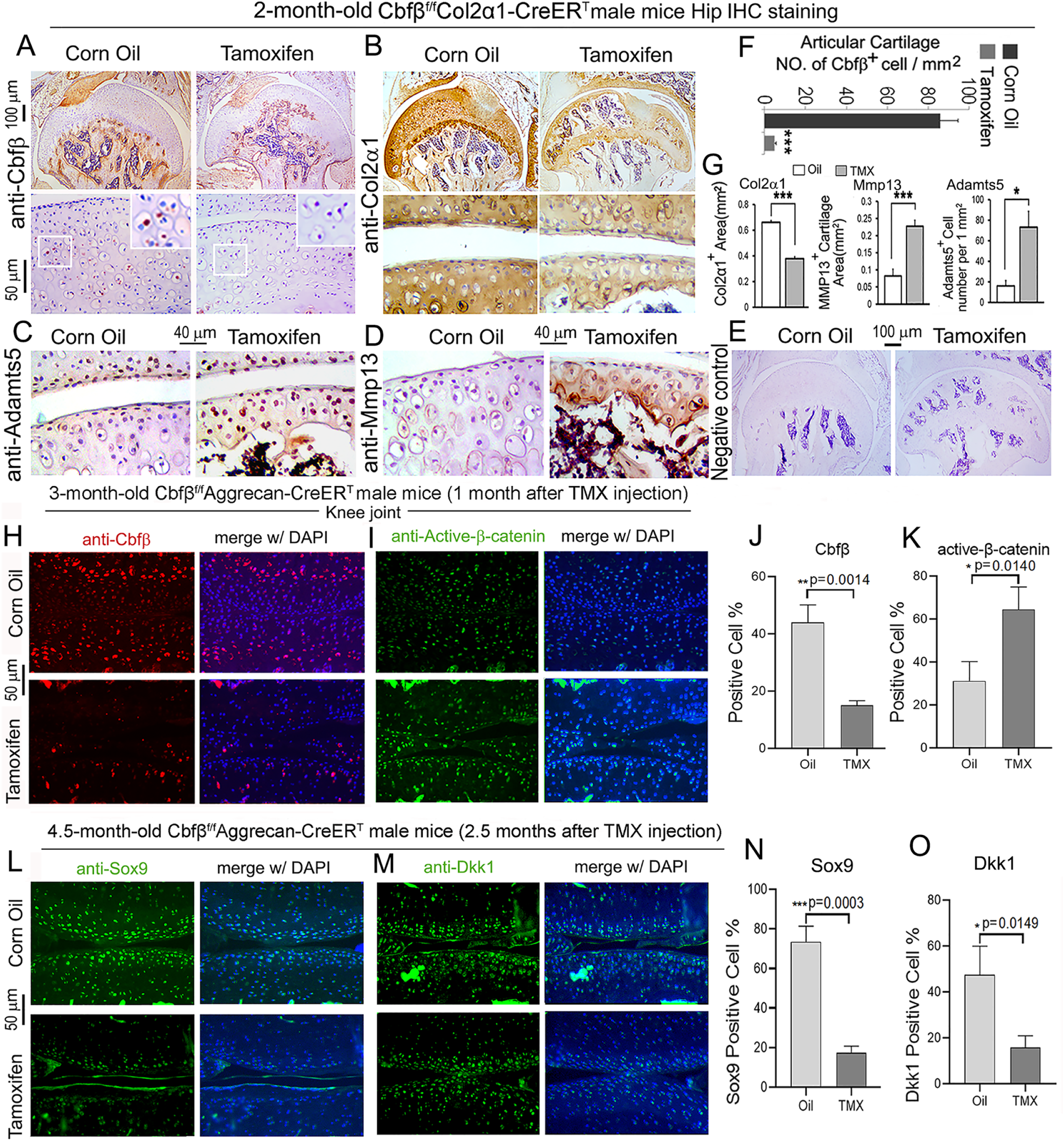
Postnatal *Cbfβ* deficiency in cartilage resulted in increased Wnt signaling, inflammatory genes expression, decreased cartilage formation genes expression in the knee articulate cartilage. **(A-E)** IHC staining of **(A)** anti-*Cbfβ,* (B) anti-Col2α1, **(C)** anti-Adamts5, and **(D)** anti-Mmp13 of hip joint from 2-month-old male *Cbfβ^f/f^Col2α1-CreER^T^* mice. **(E)** Negative control of (A-D). **(F)** Quantification for (A). **(G)** Quantification for (B-D). **(H-I)** IF staining of (**H**) anti-*Cbfβ* and **(I)** Active-β-catenin of knee joint from 3-month-old male *Cbfβ^f/f^Aggrecan-CreER^T^* mice. **(J-K)** Quantification of (H) and (I). **(L-M)** IF staining of **(L)** anti-Sox9, and **(M)** anti-Dkk1 of knee joint from 4.5-month-old male *Cbfβ^f/f^Aggrecan-CreER^T^* mice with oil injection or TMX injection. **(N-O)** Quantification of (L) and (M). Data are shown as mean ± SD. n= 3. *p < 0.05, **p < 0.01, ***p < 0.001.

As previous data had shown that *Cbfβ*-deficiency impaired articular cartilage regeneration, signaling pathways that regulate OA was our next focus, such as Wnt signaling. IF staining showed efficient Cbfβ deletion in mouse articular cartilage (**Fig. 6H, J**). Moreover, IF staining of active β-catenin showed that in knee joint articular cartilage of *Cbfβ^f/f^Aggrecan-CreER^T^* mice, there is increased Active-β-catenin expression compared to control (**Fig. 6I, K**). This confirmed that *Cbfβ* has an important role in regulating Wnt/β-catenin signaling pathway. IF staining of 4.5-month-old oil-injected *Cbfβ^f/f^Aggrecan-CreER^T^* and TMX-injected *Cbfβ^f/f^Aggrecan-CreER^T^*mice knee joints articular cartilage further exhibited that Sox9 protein expression was decreased in the Cbfβ deficient joint (**Fig. 6L, N**). As Sox9 is involved in articular cartilage formation, this observation suggests that *Cbfβ* is involved in regulating articular cartilage formation (**Fig. 6L, N**). Expression of Dickkopf-1 (Dkk1), a Wnt signaling inhibitor, was decreased expression in knee joints articular cartilage of 4.5-month-old *Cbfβ^f/f^Aggrecan-CreER^T^* mice, indicating that *Cbfβ* plays a role in regulating articular cartilage homeostasis through the Wnt signaling pathway by inhibiting Dkk1 (**Fig. 6M, O**).

### Locally administrated AAV-mediated *Cbfβ* overexpression inhibited β-Catenin expression and enhanced Yap expression in knee joints articular cartilage of ACLT-induced OA mice

To further characterize the mechanism underlying *Cbfβ* regulates articular cartilage in both physiological conditions and pathological conditions, we applied locally administrated AAV-mediated *Cbfβ* overexpression as a Gain-of-Function approach. We first proved locally administrated AAV can successfully infiltrate knee joints articular cartilage by using AAV-luc-YFP infection in mice **(SFig.1).** We then analyzed β**-**Catenin expression and Yap expression at the knee joints articular cartilage of 6.5-month-old WT mice with ACLT surgery that were either administered AAV-YFP as control or AAV-*Cbfβ* by intra-articular injection (**Fig. 7**). AAV-mediated *Cbfβ* overexpression decreased about 2.5-fold Active-β-catenin expression at the knee joints articular cartilage compared to AAV-YFP ACLT group (**Fig. 7A, B, C, H**). AAV-mediated *Cbfβ* overexpression increased Yap expression about 3.5-fold in the ACLT knee joints articular cartilage compared to the AAV-YFP ACLT group (**Fig. 7D, E, F, I**). These results from the Gain-of-Function approach confirmed that Cbfβ regulates Wnt/β-catenin and Hippo/Yap signaling pathways in articular cartilage homeostasis and suggests that local over-expression of *Cbfβ* could be an effective target for OA treatment.

**Figure 7.**
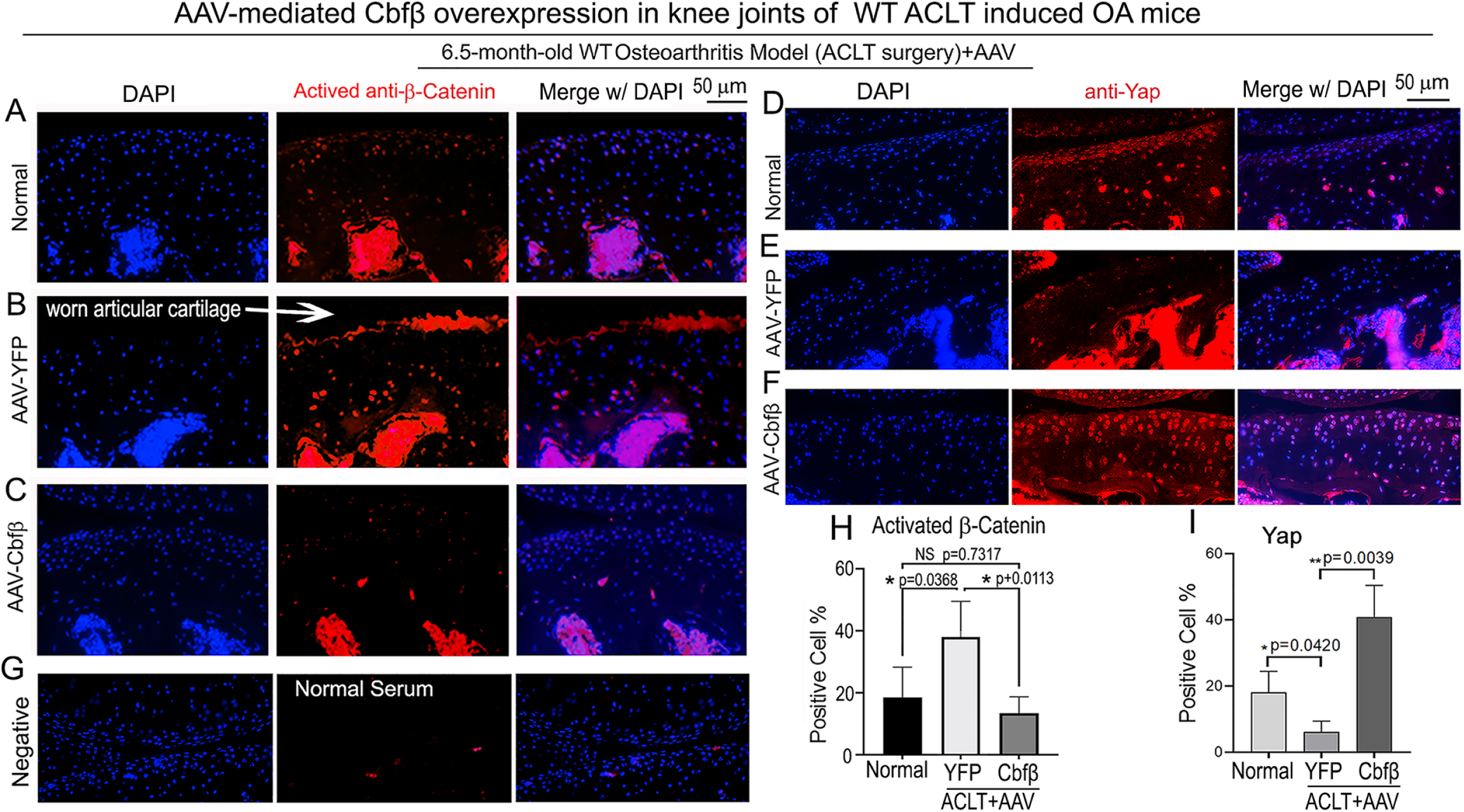
Locally administrated AAV-mediated *Cbfβ* overexpression inhibited β-Catenin expression and enhanced Yap expression in knee joints articular cartilage of ACLT-induced OA mice. **(A-C)** IF staining of anti-active-β-catenin in the knee joints articular cartilage of 6.5-month-old male **(A)** Normal WT, **(B)** AAV-YFP with ACLT surgery, and **(C)** AAV-*Cbfβ* mice with ACLT surgery (n=3). **(D-F)** IF staining of anti-YAP in the knee joints articular cartilage of 6.5-month-old **(D)**, **(E)** AAV-YFP ACLT surgery, and **(F)** AAV-*Cbfβ* mice with ACLT surgery (n=3). **(G)** Negative control of (**A-F**). **(H)** Quantification of (A-C). **(I)** Quantification of (D-F). Data are shown as mean ± SD.

### Deficiency of *Cbfβ* decreased the expression of *Yap*, and Smad2/3 and increased Mmp13 expression, and overexpression of *Cbfβ increased* Yap expression and decreased β-catenin expression

To further explore the regulatory mechanism through *in vitro* studies, we used Alcian Blue staining of primary chondrocytes prepared from newborn *Cbfβ^f/f^Col2α1-Cre* mice growth plates and showed significantly reduced matrix deposition in mutant chondrocytes, which was reflected by weaker Alcian Blue staining of the cells on days 7, 14, and 21 (**SFig. 2A, B**). Moreover, Cbf*β* overexpression in ATDC5 (chondrocyte cell line) showed about 2-fold increased Yap protein level compared to control (**Fig. 8A, B**).

**Figure 8.**
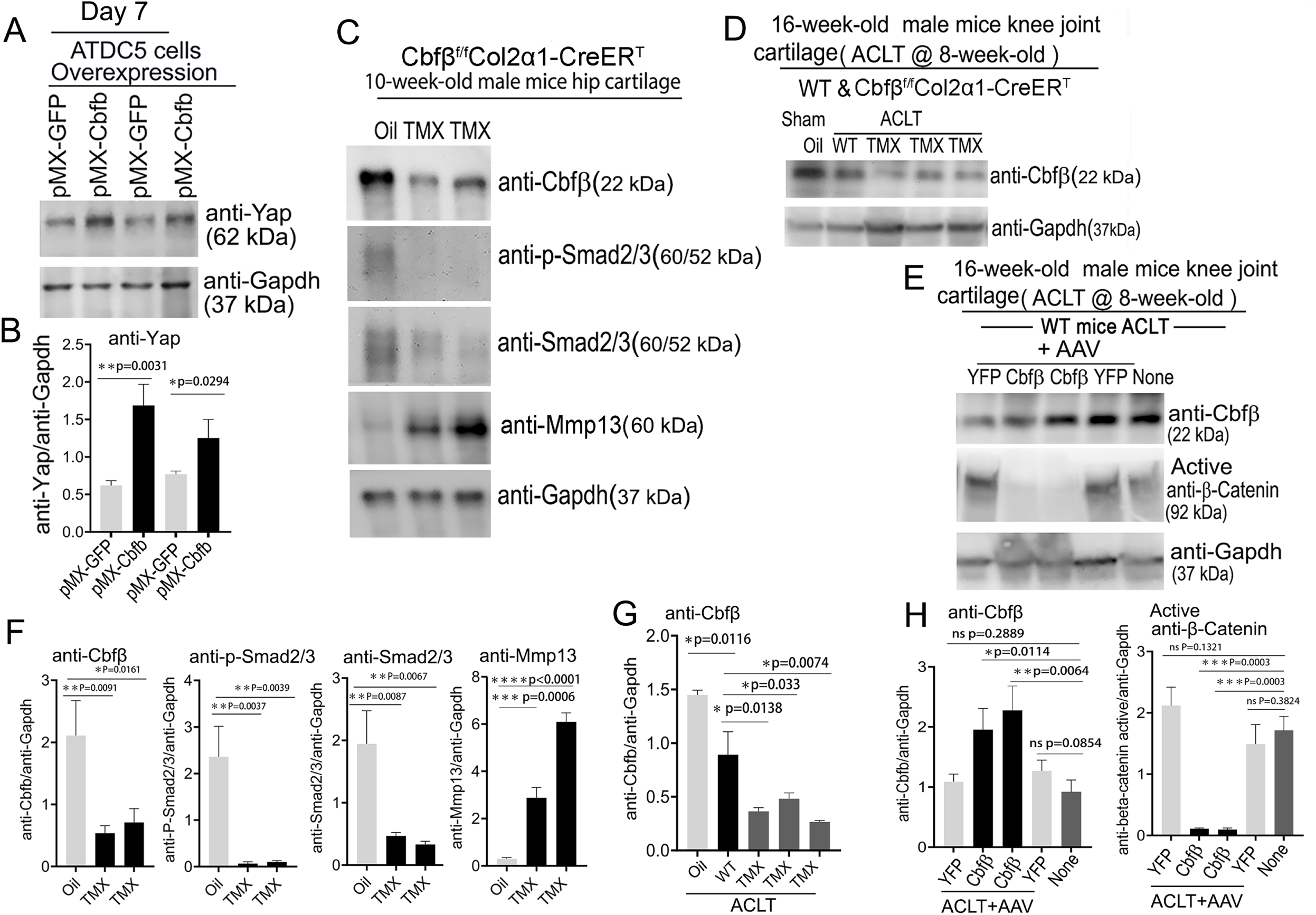
Deficiency of *Cbfβ* protein levels increased β-catenin and articular cartilage degradation markers while also reducing Yap signaling activation and *Col2α1*. **(A)** Western blot showing protein expression level of Yap in ATDC5 cells (n=3). **(B)** Quantification of Yap protein levels in (A). **(C)** Western blot of 10-week-old male hip cartilage from *Cbfβ^f/f^Col2α1- CreER^T^* mice injected with either oil or TMX showing the expression of *Cbfβ*, p-Smad2/3, Smad2/3, and Mmp13 (n=5). **(D)** Western blot of knee joint cartilage from 16-week-old male WT and *Cbfβ^f/f^Col2α1-CreER^T^*mice with ACLT surgery and injected with either oil or TMX showing the expression of *Cbfβ* (n=6). **(E)** Western blot of WT mice knee joint cartilage from 16-week-old male mice with ACLT surgery, treated with AAV-luc-YFP or AAV-*Cbfβ*, and injected with either oil or TMX showing the expression of *Cbfβ* and active β-catenin (n=6). **(F)** Quantification of (**C**). **(G)** Quantification of (**D**). **(H)** Quantification of (E). Data are shown as mean ± SD. *p < 0.05, **p < 0.01, ***p < 0.001. NS Not Significant. **Figure 8-Source data.** Raw western blot images for Figure 8A**, C-E.**

We next examined the *Cbfβ*, p-Smad2/3, Smad2/3 and Mmp13 protein level changes in hip articular cartilage in TMX injected *Cbfβ^f/f^Col2α1-CreER^T^* mice. The western blot of the hip cartilage samples showed about 3.5-fold, 10-fold and 3.5-fold decrease in protein levels of *Cbfβ*, p-Smad2/3, and Smad2/3 respectively, and a 10-fold increase in the protein level of Mmp13 (**Fig. 8C, F**). To determine whether ACLT induced OA affects *Cbfβ* protein levels, we detected *Cbfβ* protein in the knee joint articular cartilage of ACLT induced OA of 16-week-old male WT mice. The result showed that *Cbfβ* protein in WT mice with ACLT surgery decreased by about 2-fold compared to the no ACLT WT mice control, and *Cbfβ* protein was decreased by about 4-fold in *Cbfβ^f/f^Col2α1-CreER^T^* mice with ACLT compared to the control (**Fig. 8D, G**). This result indicated that low expression of *Cbfβ* may be a cause of OA pathogenesis.

To further characterize the mechanism by which *Cbfβ* regulates β-catenin expression in ACLT induced OA, the protein samples from the knee joint articular cartilage of 16-week-old male WT mice with ACLT treated with AAV-*Cbfβ* overexpression were analyzed by western blot which showed about 2-fold increased *Cbfβ* protein level in the knee joint articular cartilage of 16-week-old male WT mice with ACLT treated with AAV-*Cbfβ* mediated overexpression (**Fig. 8E, H)**, and about 10-fold decreased protein level in Active-β-catenin when compared to mice with no AAV-*Cbfβ* treatment control (AAV-YFP) (**Fig. 8E, H)**. Together, these data show that *Cbfβ* plays a central role in regulating the hip and knee joints articular cartilage homeostasis through Wnt/β-catenin, Hippo/Yap and TGFβ signaling pathways.

### Adeno-associated virus (AAV)-mediated *Cbfβ* overexpression protects against ACLT induced OA

To investigate the therapeutic effect of *Cbfβ* in ACLT induced OA, AAV-*Cbfβ* was locally administrated for AAV-mediated Cbfβ overexpression in knee joints articular cartilage of ACLT-induced OA mice. We performed X-rays and SO staining on WT mice with or without ACLT surgery and with either no treatment, AAV-YFP control treatment, or AAV-*Cbfβ* treatment (**Fig. 9**). In the X-ray images, yellow arrows indicate normal joint space; white arrows indicate worn articular cartilage; blue arrows indicate osteophytes; red arrows indicate joint space loss (**Fig. 9A B**). We observed that 22-week-old male WT mice with ACLT surgery developed an OA phenotype including unclear borders, narrow joint space, and hyperosteogeny (**Fig. 9A C**). We noticed that in the mice with ACLT surgery, the knee which was not operated on also developed a slight OA phenotype with narrow joint space. Notably, we observed that 22-week-old male WT mice with ACLT surgery treated with AAV-*Cbfβ* did not develop unclear borders, narrow joint space, hyperosteogeny, or worn articular cartilage when compared to 22-week-old male WT mice with ACLT surgery that were treated with AAV-YFP (**Fig. 9A, B**). To further investigate the role of *Cbfβ* in pathological OA through gain-of-function, we performed SO staining and histological analysis. AAV-mediated *Cbfβ* overexpression treatments were administrated to ACLT surgery-induced OA mouse models. First, we performed ACLT surgery on 8-week-old WT mice administered with AAV-YFP as control or AAV-*Cbfβ* by intra-articular injection. SO staining of the mice at 16-weeks-old showed severe articular cartilage loss in AAV-YFP treated OA mice knees, with articular cartilage degradation and osteophytes, while the AAV-*Cbfβ* treatment group had attenuated articular cartilage damage and significantly reduced OARSI scores compared to AAV-YFP control (**Fig. 9C, D, E**). These data suggest that AAV-*Cbfβ* treated mice were protected from ACLT-induced OA damage compared to the control, and that local overexpression of *Cbfβ* could be an effective therapeutic strategy for OA treatment.

**Figure 9.**
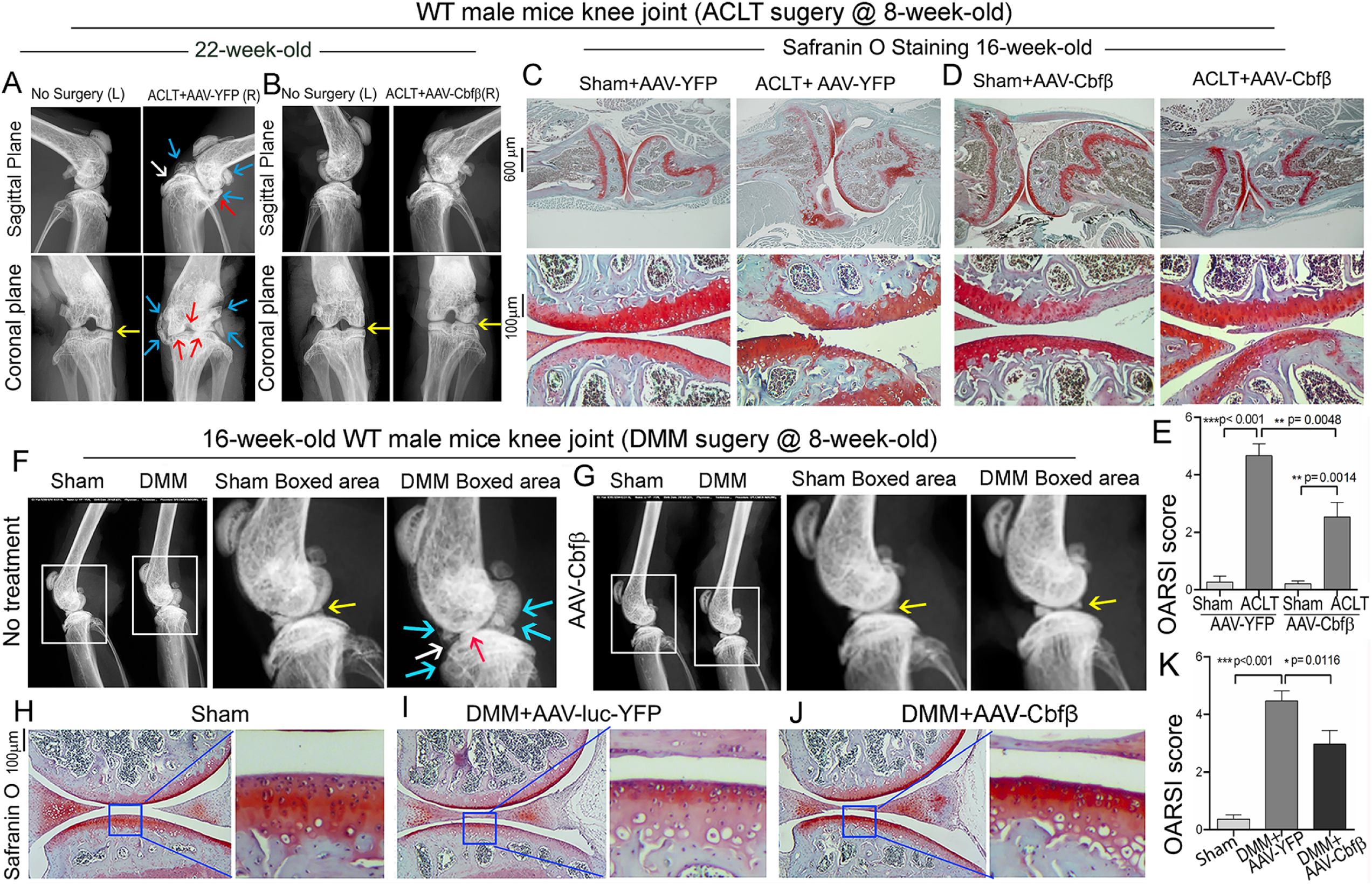
Adeno-associated virus (AAV)-mediated *Cbfβ* overexpression protects against ACLT mechanical OA. **(A-B)** X-ray images of the knee joints of 22-week-old male WT mice with ACLT surgery at 8-weeks-old with **(A)** AAV-YFP treatment and **(B)** AAV-*Cbfβ* treatment (n=15). Yellow arrows indicates normal joint space; White arrows indicate worn articular cartilage; blue arrows indicate osteophytes; red arrows indicate joint space loss. **(C-D)** SO staining of knees from 16-week-old male WT mice with (C) AAV-YFP (control) or (D) AAV-Cbfβ treatment in ACLT mediated OA (ACLT surgery at 8-weeks-old) (n=5). **(E)** Knee joint of OARSI score of (C) and (D). **(F-G)** X-ray images of mouse knee joints of 16-week-old male mice after sham/DMM surgery with **(F)** no treatment or **(G)** AAV-*Cbfβ* treatment (n=15). White arrows: osteophytes and worn articular cartilage. (**H-J**) SO staining of knee joints of 16-week-old mice after sham/DMM surgery (DMM surgery at 8-weeks-old) with (H) Sham no treatment, (I) DMM surgery AAV-YFP treatment, or (J) AAV-*Cbfβ* treatment (n=5). **(K)** Knee joint OARSI score of (H-J). The results are presented as the mean ± SD, *p < 0.05, **p < 0.01, ***p < 0.001. DMM surgery AAV-YFP treatment group shows severe cartilage damage, osteophytes, and delocalized knee joint, while the AAV-*Cbfβ* treated group shows less cartilage loss and osteophytes than control.

We also used the surgical destabilization of the medial meniscus (DMM) surgery-induced OA model to test *Cbfβ’s* role in protecting against OA. We observed that In DMM surgery-induced OA. The OA phenotype was evident (**Fig. 9F, G**) as indicated by blue and white arrows. Notably, in DMM surgery-induced OA with AAV-Cbfβ treatment the OA phenotype was not seen, and only slightly increased subchondral bone density was observed **(Fig. 9F, G)**. To further investigate the role of *Cbfβ* in pathological OA through a gain-of-function approach, AAV-mediated *Cbfβ* overexpression treatments were administrated to DMM surgery-induced OA mouse models. SO staining of the mice at 16-weeks-old showed articular cartilage loss in AAV-YFP treated OA mice knees, with degraded articular cartilage and osteophytes, while the AAV-*Cbfβ* treatment group displayed attenuated articular cartilage damage and significantly reduced OARSI scores compared to AAV-YFP control (**Fig. 9G, H, I, J, K**). Consistent with our SO staining of ACLT model of OA, evaluation of mice knee joints in DMM-induced OA showed loss of articular cartilage, decreased joint space, and increased OARSI score (**Fig. 9H, I, K**). Compared to AAV-YFP controls, treatment with AAV-*Cbfβ* attenuated articular cartilage damage and significantly reduced OARSI scores compared to AAV-YFP control **(Fig. 9I-K).** Thus, local overexpression of Cbfβ could be a novel and effective target for the treatment of osteoarthritis.

## DISCUSSION

In our current study, we showed that the deletion of *Cbfβ* in postnatal mice cartilage caused severe spontaneous OA through *Cbfβ’s* regulation of multiple key signaling pathways. We demonstrated that the changes of OA-related gene expression in the articular cartilage of aging-associated and *Cbfβ* deficiency induced OA included downregulated Sox9, Dkk1, Yap, and p-Smad2/3, and upregulated Wnt5a and Wnt/β-catenin. We conclude that *Cbfβ* reverses articular cartilage regeneration and repair by modulating multiple key signaling pathways, including the Wnt/β-catenin, TGF-β, and Hippo/YAP pathways.

Our previous studies have proven *Cbfβ’s* important role in bone skeletal development(16, 19). *Cbfβ and Runx1* plays crucial roles in regulating both chondrocytes and osteoblast in bone(15, 49–52). *Cbfβ* is known to bind to Runx proteins (*Runx1*, *Runx2*, *Runx3*) through the Runt domain, and exon 5 of the *Cbfβ* gene is essential for *Cbfβ*-Runx binding ability(20). Recent studies revealed that *Cbfβ* played an important role in stabilizing Runx proteins(20, 21). In this study, we found *Runx2* reduced expression in *Cbfβ*-deficient OA hip articular cartilage as shown by heatmap analysis. Several OA susceptibility genes were identified through a genome-wide DNA methylation study in OA cartilage tissue, including *Runx1* and *Runx2*(53). *Runx1* was reported to be highly expressed in knee superficial zone chondrocytes, to regulate cell proliferation(54). In addition, *Runx1* expression was increased in knees with OA(54). Furthermore, *Runx1* mRNA injection showed a protective effect on surgically induced OA knees(5). In our study, we found *Runx1* expression in the superficial zone and columnar chondrocytes of hip cartilage. In *Cbfβ* deletion mice, *Runx1* expression was largely decreased at both the mRNA and protein level, which could directly relate to OA development. In contrast, *Runx2* is known to promote OA formation by upregulating Mmp13(43, 55). However, a recent publication has shown that overexpression of *Runx2* driven by the ColX promoter in mice showed delayed chondrocyte maturation and decreased susceptibility to develop OA, indicating that temporally and spatially different expressions of *Runx2* may play opposite roles in OA(56).

Our data indicated that *Cbfβ* deletion upregulated the Wnt canonical signaling pathway during chondrocyte homeostasis. RNA-seq analysis showed increased expression of Wnt10b, Wnt2b, the activator of Wnt signaling, and decreased Wnt antagonistic inhibitor Sost expression in *Cbfβ^f/f^Col2α1-CreER^T^*mice hip tissue. The Wnt signaling pathway has been implicated in OA in both clinical data and animal models(57). Wnt/β-catenin pathway inhibitors DKK1, Axin2, and alternative Wnt ligand Wnt5a, were highly expressed in human OA samples(58–60). Meanwhile, Gremlin 1 (Wnt signaling antagonists), Frizzled-related protein (Wnt receptor), and DKK1 are recognized as key regulators of human articular cartilage homeostasis(61). Furthermore, functional variants within the secreted frizzled-related protein 3 gene (Wnt receptor) are associated with hip OA in females(62). These data indicate that Wnt signaling is closely related to human OA formation. In experimental mouse models, both repression(63) and forced activation(64) of β-catenin caused OA. Yet how canonical Wnt signaling is dysregulated in OA remains unclear. Our data demonstrate that *Cbfβ* enhances articular cartilage regeneration and repair by orchestrating multiple key signaling pathways, including Wnt/β-catenin. Our results provide new insights into how Wnt canonical signaling is regulated during OA pathogenesis which may lead to novel therapies for the treatment of degenerative joint diseases.

Our study supports that *Cbfβ* promotes the Hippo/YAP pathway in chondrocyte homeostasis. YAP has been reported to upregulate chondroprogenitor cells proliferation and inhibit chondrocyte maturation(65), and Yap1 and *Runx2* protein-protein interaction has been previously confirmed(65). In our studies, we found that in *Cbfβ*-deficient cartilage, YAP transcriptional target genes Wnt5a/b were highly decreased. Thus, *Cbfβ* may promote YAP expression by regulating *Runx2* expression and *Cbfβ/Runx2*-YAP protein-protein interaction. Some research has also indicated that Wnt5a/b-YAP signaling antagonizes canonical Wnt/β-catenin signaling and decreases expression of a panel of the major β-catenin/TCF target genes(66). However, further study is needed. Our results demonstrate that the high expression level of YAP in cartilage suppresses Wnt/β-catenin pathways, thus leading to the OA phenotype seen in aging mice.

TGF-β signaling also plays key roles in the development of the spontaneous OA phenotype as shown by our data. Maintaining homeostasis in articular cartilage and subchondral bone requires precise control of the TGF-β signaling pathway(7, 24). However, various components of the TGF-β signaling pathway, along with *Cbfβ*, have been shown to decrease with age, illustrating a possible mechanism in the development of OA(21). *Cbfβ* and *Runx1* have been revealed to be mediators of TGF-β signaling, with the activation of TGF-β signaling having been shown to increase *Cbfβ* and *Runx1* expression and *Cbfβ/Runx1* heterodimer formation, while *Cbfβ* deletion attenuates TGF-β signaling (24). Our RNA sequencing results illustrate a similar pattern, where *Cbfβ* conditional knockout resulted in concomitant reduction of TGF-β1 expression in cartilage cells. The previously cited paper also reported that the disruption of TGF-β signaling by the deletion of *Cbfβ* in articular chondrocytes showed an increase in catabolic cytokines and enzymes interleukins and matrix metalloproteinases (24). Furthermore, elevated levels of TGF-β1 in subchondral bone has been linked to the pathogenesis of OA(7). Our work also indicated that *Cbfβ* conditional knockout within *Cbfβ^f/f^Aggrecan-CreER^T^*after TMX induction resulted in significantly elevated Mmp13, suggesting a possible therapeutic target for the prevention or reduction in OA progression.

Another important component of TGF-β signaling is Smad proteins, which are required to be phosphorylated in order to facilitate the transcription of bone and cartilage homeostasis mediators(67). Our RNA sequencing and western blot results demonstrated both altered expression and activation of several other SMADs and components of the TGF-β signaling pathways such as Smad2 and Smad3. The previous cited study shows contradictory results to this study, showing elevated p-Smad3 protein in *Cbfβ* conditional knockout mice knee joint articular cartilage(24). Such difference might be due to various factors. Firstly, TGF-β signaling has been reported for both protective and catabolic roles in the pathogenesis of OA(68). In fact, study has shown that short and long stimulation of TGF-β has completely opposite effects on the cartilage health(69). The dual role of TGF-β signaling might cause the discrepancies observed in downstream regulators. Additionally, TGF-β signaling is complex in the way that many signaling pathways can affect it signaling. For example, differed inflammatory states results in altered downstream TGF-β signaling, which can also be an explanation for the deviance in results(70). As such, the mechanism by which *Cbfβ* expression affects p-Smad2/3 requires further elucidation as it is unclear whether this functions by either a positive or negative feedback mechanism. Nevertheless, our study has well proved that *Cbfβ* has a crucial function in maintaining TGF-β signaling and chondrocyte homeostasis.

In addition to those important pathways mentioned above, we also identified several significantly differentially upregulated/downregulated genes in *Cbfβ* conditional knockout mice, through volcano plot analysis of RNA-seq data. This includes decreased expression of Fabp3, Nmrk2, Csf3r, Rgs9, Plin5, Rn7sk, Eif3j2 and increased expression of Cyp2e1, Slc15a2, Alas2, Hba-a2, Lyve1, Snca, Serpina1b, Hbb-b1, Rsad2, Retn, and Trim10 in *Cbfβ* conditional knockout mice. We here discussed the possible regulatory role of *Cbfβ* in Fabp3, Plin5, and Rsad2. However, other differentially expressed genes that have not been discussed could also have potential important role in understanding the mechanism of *Cbfβ* in regulating chondrocyte homeostasis in OA pathogenesis. Those genes therefore need to be further studied.

In summary, we found that *Cbfβ* deletion in postnatal cartilage caused severe OA through the dysregulation of Wnt signaling pathways and overexpression of *Cbfβ* protects against OA. Our study notably revealed that *Cbfβ* is a key transcription factor in articular cartilage homeostasis and promotes articular cartilage regeneration and repair in OA by orchestrating Hippo/YAP, TGF-β, and Wnt/β-catenin signaling. The novel mechanism provides us with more insights into OA pathogenesis while also providing potential avenues for OA treatment and prevention.

## MATERIALS AND METHODS

### Generation of Cbfβ inducible CKO mice

The *Cbfβ^f/f^* (Stock No: 008765) and *Aggrecan-CreER^T^*(Stock No: 019148) mouse lines were purchased from Jackson Laboratory. *Col2α1-CreER^T^* mice line was generated and kindly provided by Dr. Di Chen(71). *Cbfβ^f/f^* mice were crossed with either *Aggrecan-CreER^T^* or Col*2α1-CreER^T^* mice to generate *Cbfβ^f/+^Col2α1-CreER^T^* or *Cbfβ^f/+^Aggrecan-CreER^T^* mice, which were then intercrossed to obtain homozygous inducible CKO (*Cbfβ^f/f^Col2α1-CreER^T^*and *Cbfβ^f/f^Aggrecan-CreER^T^*) mice. The genotypes of the mice were determined by polymerase chain reaction (PCR). Both male and female mice of each strain were randomly selected into groups of five animals each. The investigators were blinded during allocation, animal handling, and endpoint measurements. All mice were maintained in groups of 5 mice with singular sex/Breeding trios (1 male:2 females) under a 12-hour light–dark cycle with ad libitum access to regular food and water at the University of Alabama at Birmingham (UAB) Animal Facility. TMX (T5648, Sigma) was dissolved in vehicle-corn oil (C8267, Sigma) in the concentration of 10 mg/ml and vortexed until clear. The solution was aliquoted and stored at -20°C. Before use, the TMX solution was warmed at 37 ℃ for 5 minutes. 2-week-old *Cbfβ*^f*/f*^ mice and *Cbfβ^f/f^Col2α1-CreER^T^* mice 8-week-old *Cbfβ^f/f^Aggrecan-CreER^T^*) mice received either TMX or corn oil by intraperitoneal (I.P.) injection continuously for 5 days (75 mg tamoxifen/kg body weight per day).

### DMM or ACLT surgery induced OA and AAV-Cbfβ transduction

8-week-old C57BL/6 wild type mice of both sexes received either ACLT surgery, DMM surgery, or sham surgery on the right knee. We administrated AAV-CMV-*Cbfβ* in a site-specific manner as described in a previous study but with minor modifications(72). Briefly, mouse *Cbfβ* cDNA (isoform 1, BC026749) was cloned into pAAV-MCS vector, which was followed by AAV transfection by the Ca2+- phosphate/DNA co-precipitation method. AAV titer was tested by the qPCR method. The right knee capsules were locally injected with 10 μl AAV-YFP or AAV-Runx1 (titer >10^10^/ml) three times on day 7, day 14, day 21 at the knee joint cavity, and euthanized 8 weeks or 10 weeks after surgery to obtain ACLT knee joint samples as described (50). Mice were harvested for X-ray and histological analysis.

### Histology and tissue preparation

Histology and tissue preparation were performed as described previously(73). Briefly, mice were euthanized, skinned, and fixed in 4% paraformaldehyde overnight. Samples were then washed with water, dehydrated in 50% ethanol, 70% ethanol solution and then decalcified in 10% EDTA for 4 weeks. For paraffin sections, samples were dehydrated in ethanol, cleared in xylene, embedded in paraffin, and sectioned at 5 μm with a Leica microtome and mounted on frosted microscope slides (Med supply partners). H&E and SO staining were performed as described previously(74). ALP staining and TRAP staining were performed with kits from Sigma.

### Radiography

Radiographs of inducible *Cbfβ^f/f^Col2α1-CreER^T^*mice were detected by the Faxitron Model MX-20 at 26 kV in the UAB Small Animal Bone Phenotyping Core associated with the Center for Metabolic Bone Disease.

### Immunohistochemistry and Immunofluorescence analysis

The following primary antibodies were used: mouse-anti-*Cbfβ* (Santa Cruz Biotechnology Cat# sc-56751, RRID:AB_781871), mouse-anti-Col2α1 (Santa Cruz Biotechnology Cat# sc-52658, RRID:AB_2082344), rabbit-anti-MMP13 (Abcam Cat# ab39012, RRID:AB_776416), rabbit-anti-ADAMTS5 (Santa Cruz Biotechnology Cat# sc-83186, RRID:AB_2242253), rabbit-anti-Sox9 (Santa Cruz Biotechnology Cat# sc-20095, RRID:AB_661282), rabbit-anti-Yap (Santa Cruz Biotechnology Cat# sc-15407, RRID:AB_2273277), rabbit-anti-Dkk1 (Cell Signaling Technology Cat# 48367, RRID:AB_2799337), and mouse-anti-Active-β-catenin(Millipore Cat# 05-665, RRID:AB_309887). Imaging was done with a Leica DMLB Microscope and a Leica D3000 fluorescent microscope and were quantified by Image J software.

### Protein sample preparation

Mouse femoral hip articular cartilage or mouse knee cartilage was isolated, washed with sterile ice cold 1x PBS twice, added with appropriate amount of 1x SDS protein lysis buffer and protease inhibitor cocktail in 1.5 ml tube. Keeping on ice, femoral hip or knee tissue were quickly cut into small pieces using small scissors in 1.5.ml tube. Centrifugation was performed at room temperature at 16,000 rpm for 30 seconds. The supernatant was then transferred to a new, pre-chilled 1.5ml centrifuge tube, discarding bone debris, and then boiled in water for 10 minutes and kept on ice. Samples were either used directly for western blot or stored at -80°C.

### Western blot analysis

Proteins were loaded on SDS-PAGE and electro-transferred on nitrocellulose membranes. Immunoblotting was performed according to the manufacturer’s instructions. The following primary antibodies were used: mouse-anti-*Cbfβ* (Santa Cruz Biotechnology Cat# sc-56751, RRID:AB_781871), rabbit-anti-MMP13 (Abcam Cat# ab39012, RRID:AB_776416), rabbit-anti-Yap (Santa Cruz Biotechnology Cat# sc-15407, RRID:AB_2273277), mouse-anti-GAPDH (Santa Cruz Biotechnology Cat# sc-365062, RRID:AB_10847862), mouse-anti-Active-β-catenin(Millipore Cat# 05-665, RRID:AB_309887), rabbit-anti-Smad3(Cell Signaling Technology Cat# 9513, RRID:AB_2286450), and rabbit-anti-pSmad3 (Cell Signaling Technology Cat# 9520 (also 9520S, 9520P), RRID:AB_2193207). Secondary antibodies were goat anti-rabbit IgG-HRP (Santa Cruz Biotechnology Cat# sc-2004, RRID:AB_631746), and rabbit anti-mouse IgG-HRP (Santa Cruz Biotechnology Cat# sc-358917, RRID:AB_10989253). Quantification of Western blot area was performed by ImageJ.

### Primary chondrocyte culture and ATDC5 cell transfection

We isolated and cultured primary chondrocytes from neonatal f/f and Cbfβ^f/f^Col2ɑ1-Cre mice as described(75). Primary mouse chondrocytes were induced for 7 days. Alcian blue staining was carried out to detect chondrocyte matrix deposition as previously described (76). We used pMXs-GFP and pMXs-3xFlag-Cbfβ (pMX-Cbfb) retroviral vectors to package and collect retroviruses, which infected ATDC5 (ECACC Cat# 99072806, RRID:CVCL_3894) cells to enhance the expression of Cbfβ. The infected ATDC5 cell line cells were induced for 7 days before harvest for protein Western blot analysis.

### Published Data Analysis

Human patient information from OA cartilage samples came from prior work for RNA-seq of knee OA compared to normal controls (Accession# GSE114007)(77) and for methylation chip comparison of hip OA compared to hip fracture controls (Accession# GSE63695)(27). Analysis and comparison were performed using GEO2R and GEOprofiles. Statistical significance was assessed using Student’s t-test. Values were considered statistically significant at p<0.05.

### RNA-Sequencing Analysis

Total RNA was isolated using TRIzol reagent (Invitrogen Corp., Carlsbad, CA) from hip articular cartilage or mouse knee cartilage and was submitted to Admera Health (South Plainsfield, NJ), who assessed sample quality with the Agilent Bioanalyzer and prepared the library using the NEBnext Ultra RNA - Poly-A kit. Libraries were analyzed using Illumina next generation sequencing and relative quantification was provided by Admera Health. Sequence reads were aligned to GRCm39/mm39 reference genome using STAR (v.2.7.9) and visualized using Integrative genomics viewer (igv v.2.16.2). Read counts were subjected to paired differential expression analysis using the R package DESeq2. Top GO downregulated categories were selected according to the *P*-values and enrichment score and illustrated as number of genes downregulated in respective category.

### Data Availability

The RNA-seq data has been deposited in the Gene Expression Omnibus (GEO) under accession code GSE253210.

### Statistical Analysis

The number of animals used in this study was determined in accordance with power analysis and our previous studies (78). In brief, our study used five mice per group per experiment. Data are presented as mean ± SD (n ≥ 3). Statistical significance was assessed using Student’s t test. Values were considered statistically significant at P < 0.05. Results are representative of at least three individual experiments. Figures are representative of the data.

### Ethics approval

All animal experimentation was approved by the IACUC at the University of Alabama at Birmingham and was carried out according to the legal requirements of the Association for Assessment and Accreditation of the Laboratory Animal Care International and the University of Alabama at Birmingham Institutional Animal Care and Use Committee. All studies follows NIH guidelines.

## Acknowledgements

This work was supported by the National Institutes of Health [AR-070135 and AG-056438 to W.C., and AR075735 and AR074954 to Y.P.L].

## Author Contributions

Study design: WC and YPL. Study conduct: WC, YL, YZ, JW, AM, YC, SZ, GZ and YPL. Data collection and analysis: WC, YL, YZ, JW, AM, YC, SZ, GZ, YLu, JZ, MM, and YPL. Drafting manuscript: WC, YL, YZ, JW, AM, YC, SZ, MM and YPL. Revising manuscript: WC, YL, YZ, JW, AM, YC, SZ, GZ, YLu, JZ, MM and YPL. All authors approved the final version of the manuscript for submission. WC and YPL take responsibility for the integrity of the data analysis.

## Abbreviations

The abbreviations used are: OA, osteoarthritis; *Cbfβ*, Core binding factor subunit β; RUNX1, Runt-related transcription factor 1; TRAP, Tartrate-resistant acid phosphatase; ALP, alkaline phosphatase, H&E, hematoxylin and eosin; SO, Safranin O; IF, immunofluorescence; OARSI, Osteoarthritis Research Society International; TMX, tamoxifen; AAV, adeno-associated virus; ACLT, Anterior cruciate ligament transection; DMM, destabilization of the medial meniscus.

## SUPPLEMENTAL FIGURES AND FIGURE LEGENDS

**Supplementary Figure 1.**
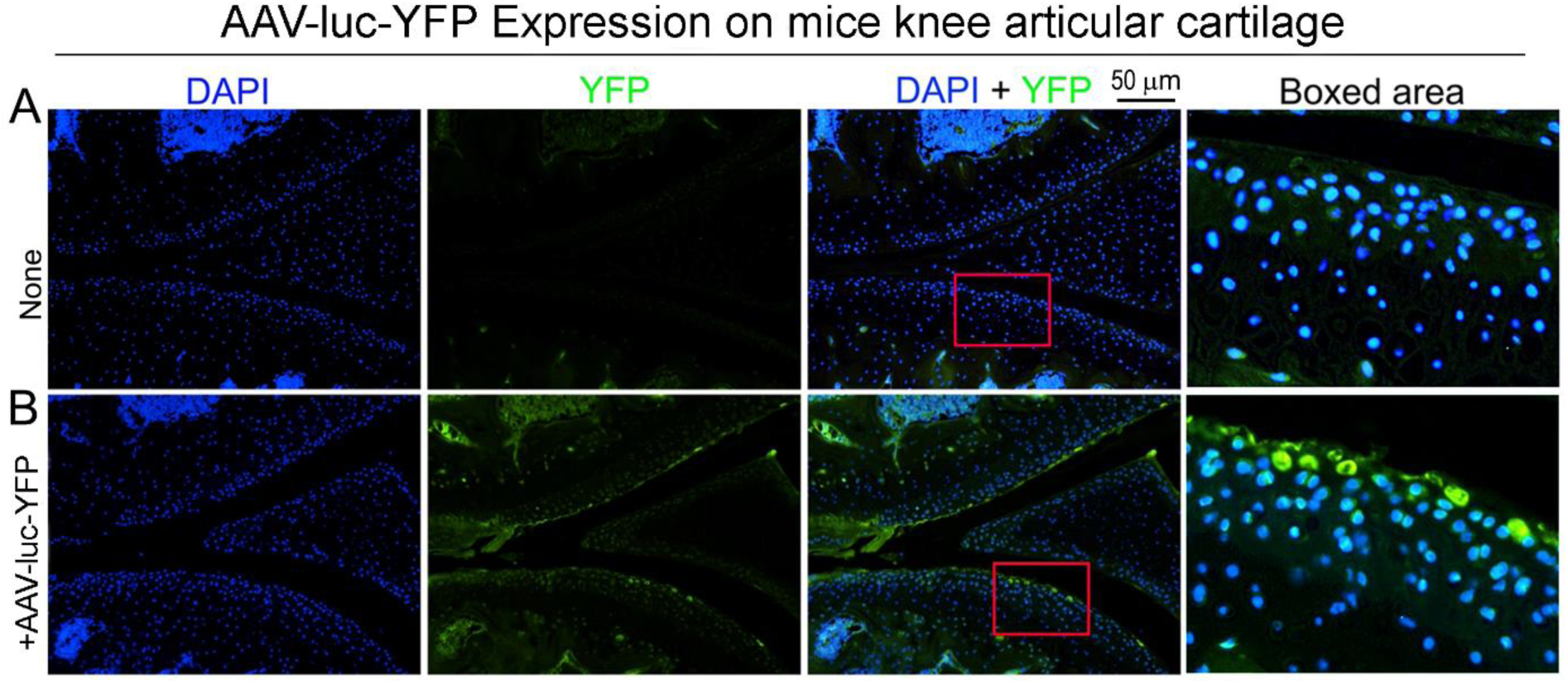
Successful AAV-luc-YFP infection in mice. Observed by fluorescence microscope**. (A)** DAPI staining for 8-weeks-old male mouse knee articular cartilage Observed by fluorescence microscope. **(B)** DAPI staining and expression for YFP in 8-weeks-old male mice knee articular cartilage Observed by fluorescence microscope. Scale bar: 50 μm (A, B). (n=3)

**Supplementary Figure 2.**
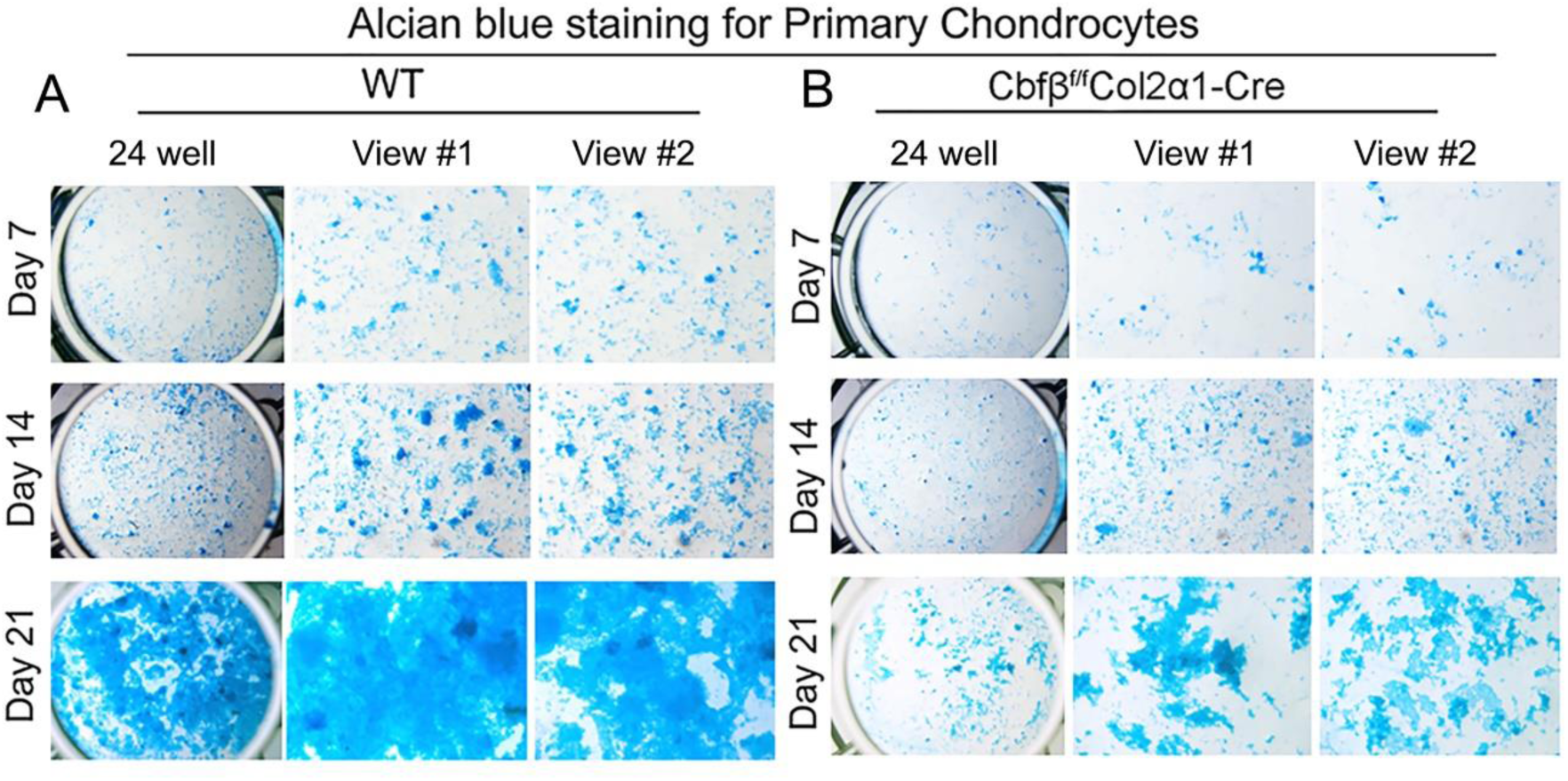
Alcian Blue staining of primary chondrocytes from *Cbfβ* deficient newborn mice show reduced matrix deposition. **(A)** Alcian Blue staining of newborn WT mouse primary chondrocytes. **(B)** Alcian Blue staining of newborn (P0) *Cbfβ^f/f^Col2α1-Cre* mouse primary chondrocytes. (n=5)

